# Locally stimulating cell migration in living tissues drives long range collective motion through cell-cell adhesion leading to accelerated migration, healing, and growth

**DOI:** 10.1101/2025.11.19.689141

**Authors:** Jeremy S. Yodh, Yubin Lin, Sumit Sinha, Vishaal Krishnan, L. Mahadevan, Daniel J. Cohen

## Abstract

Collective cell migration is critical in a range of biophysical processes spanning wound healing to tumor metastasis. It is therefore important to develop techniques for regulating cell motion, but while we have developed powerful migration tools from optogenetics and bioelectricity, the complex mechanics of collective systems make it difficult to determine where and when to apply these stimuli. For example, here we begin with a circular sheet of skin cells with a central hole, or “wound”, and show that globally stimulating all cells to migrate radially inwards to close the gap through electrotaxis causes catastrophic mechanical damage, emerging from strong cell-cell adhesion, which we explain with an active elasticity model. We propose a solution inspired by sheepherding based on using local stimulation to learn the collective perturbation-response of the group and then drive tissue motion. First, we induce local electrotaxis to characterize the impulse-response function of quasi-1D strips of skin tissue, discovering that hyper-local stimulation triggers long-range tissue response over a length scale set by cell-cell adhesion also predicted by our model. Based on this, we apply local, concentric ring fields to in vitro circular wounds. Continuously driving cells towards the wound core kinetically traps the tissue in a jammed state and freezes healing. Pulsing the stimuli allows the tissue to relax and fluidize again, accelerating migration. Finally, we integrate the key length and timescales of tissue mechanics into a biophysically-informed continuum control model. The model’s predictive framework helps determine where and when to apply stimulation for optimal tissue growth which, when tested, accelerates healing 5-fold.

---

Collective cellular motion, which coordinates and enables processes from embryogenesis to healing to cancer invasion [1–3], is among the best characterized forms of collective behavior and has become as a powerful domain for studying non-equilibrium statistical physics and crowd dynamics. Even simple monolayer sheets of cells (typically from skin, kidney, gut, blood vessels, and mammary gland [4–8]) have revealed potent and exotic physics from massive density fluctuations, activity-driven phase separation, and active instabilities (see [9–12] and references therein), because these materials are active, i.e., they can convert internal energy into net motion. Moreover, the physics of collectives have connected our understanding of branches of soft matter physics (e.g., granular media, viscoelasticity, active matter) to our understanding of cellular systems and vice versa. However, as we come to better understand the physics underlying collective cell motion, the field is advancing to a new domain centered on exploring how to leverage these physical principles to rationally drive cell and other collective systems. Here, we investigate what physical constraints set the length and timescales of how input stimuli propagate and dissipate in a living tissue, and we use the underlying physics to determine *where* and *when* to apply stimuli to a tissue to best regulate collective cell migration in a healing experiment.

Regulating collective motion within a tissue is further complicated because even small tissues are 𝒪 (10^3^−10^4^) cells *in vitro*, and the meso-scale physics of the group emerges from local interactions. Trying to address each agent individually is both computationally infeasible and ignores emergent mechanics. One obvious approach is to apply a global stimulus to all agents. However, global commands often clash with the local dynamics and endogenous collective behaviors of the group. In tissues, global commands can jeopardize tissue health (a validated strategy to circumvent damage is to override existing collective dynamics with drugs [13]). Another classic example is the panic problem where overly incentivizing all individuals to rush to the nearest exit can cause granular jamming within human crowds [14]. Even more surprisingly, globally commanding all cars on a circular track to maintain fixed speed and distance from their neighbors can trigger a traffic jam due to local instabilities in driver reaction times [15].

A bio-inspired solution is to derive the perturbation-response of the group with local, rather than global inputs. Nature exhibits fantastic examples of this through predator-prey interactions where the few drive the many. The most elegant system in this domain, sharpened by ~7000 years of practice, is sheepherding, where a single dog can drive a flock. Here, the dog iteratively applies “pressure” that adapts to the shape, size, and position of the flock, dynamically oscillating between perturbations that gather the group and those which drive it [16–18]. In essence, the sheepdog is performing a linear-response experiment with a local cue to learn the properties of the flock. Ultimately, this strategy of local spatiotemporal control is substantially more energy efficient because local perturbations can guide, but need not excite, the entire group. In fact, a way to mitigate the earlier example of spontaneous traffic jams under global rules is to introduce even just one autonomous vehicle into the group whereupon its local effects can effectively dampen traffic jamming and improve global speed and fuel efficiency [15].

Here, we explore whether we can adapt this bioinspired motif of local perturbation to global response to characterize and manipulate the motion of cellular collectives with over 10,000 cells. We chose sheets of skin cells (primary mouse keratinocytes) as the model system because their collective properties are understood to emerge from strong cell-cell adhesion, and it is experimentally possible to tune the adhesion strength, cell number density, and the tissue size and shape. Here, we engineered large circular monolayers of skin cells with central wounds, allowing us to compare natural self-healing to globally versus locally stimulated healing. This accelerated healing framework effectively probes the coupling between collective biophysics, perturbation response, and functional outcomes.

As an external stimulus, we use weak direct cur-rent (DC) electric fields, harnessing a well-characterized process called *electrotaxis*, where DC ionic currents of 𝒪(1 V*/*cm) *in vivo* typically direct cell migration along the field and play a key role in native cell migration and healing [19–29]. These fields can be mimicked and amplified *in vitro*, and applied globally or locally, allowing us to compare the collective perturbation responses to global electric stimuli versus local stimuli.

We first use this strategy to derive the perturbation response of living skin tissues for global and local stimulation. After showing that global stimulation produces mechanical damage [30] to the tissue, we perform a series of impulse-response experiments, where we characterize how far the effects of local stimuli applied only to the middle of the tissue propagate into the tissue bulk. This data, combined with continuum active elastic modeling, reveal how and why a local stimulus produces a much longer-range response and validate our local to global strategy. We next attempt to use the length scales derived from the impulse-response experiments to determine where to locally stimulate a healing tissue, encountering problems reminiscent of granular jamming and leading us to explore the relaxation timescales during stimulation. Finally, we took the physical length and timescales derived from our experiments and combined them with a physics-derived control model that served to optimize both the placement and timing of stimulation to maximize the mechanical coupling efficiency of the local inputs to long-range migration and accelerate healing of the model wound 5-fold over the native rate.

Together, this work demonstrates how characterizing the perturbation response of a living collective can generate the physical parameters and intuition needed to more effectively regulate and drive collective motion.

## RESULTS

### Global commands cause mechanical damage from collective coupling

We begin with the common biological example of wound closure, or epithelial self-healing, which underlies processes spanning dorsal and neural tube closure in development to basic puncture healing [31, 32]. Here, we engineered a sheet of skin cells with a hole in the center (Fig. 1A). This system is capable of slow, collective self-healing (Supplemental Video 1), and it would seem logical that a command (an electric field) directing cells to migrate inwards should accelerate the process. Although this global command initially triggered rapid, global inward migration, this came at the cost of catastrophic failure of the tissue at the wound margin (Supplemental Video 2), which retracted and made the wound *larger* (Fig. 1B).

**FIG. 1.**
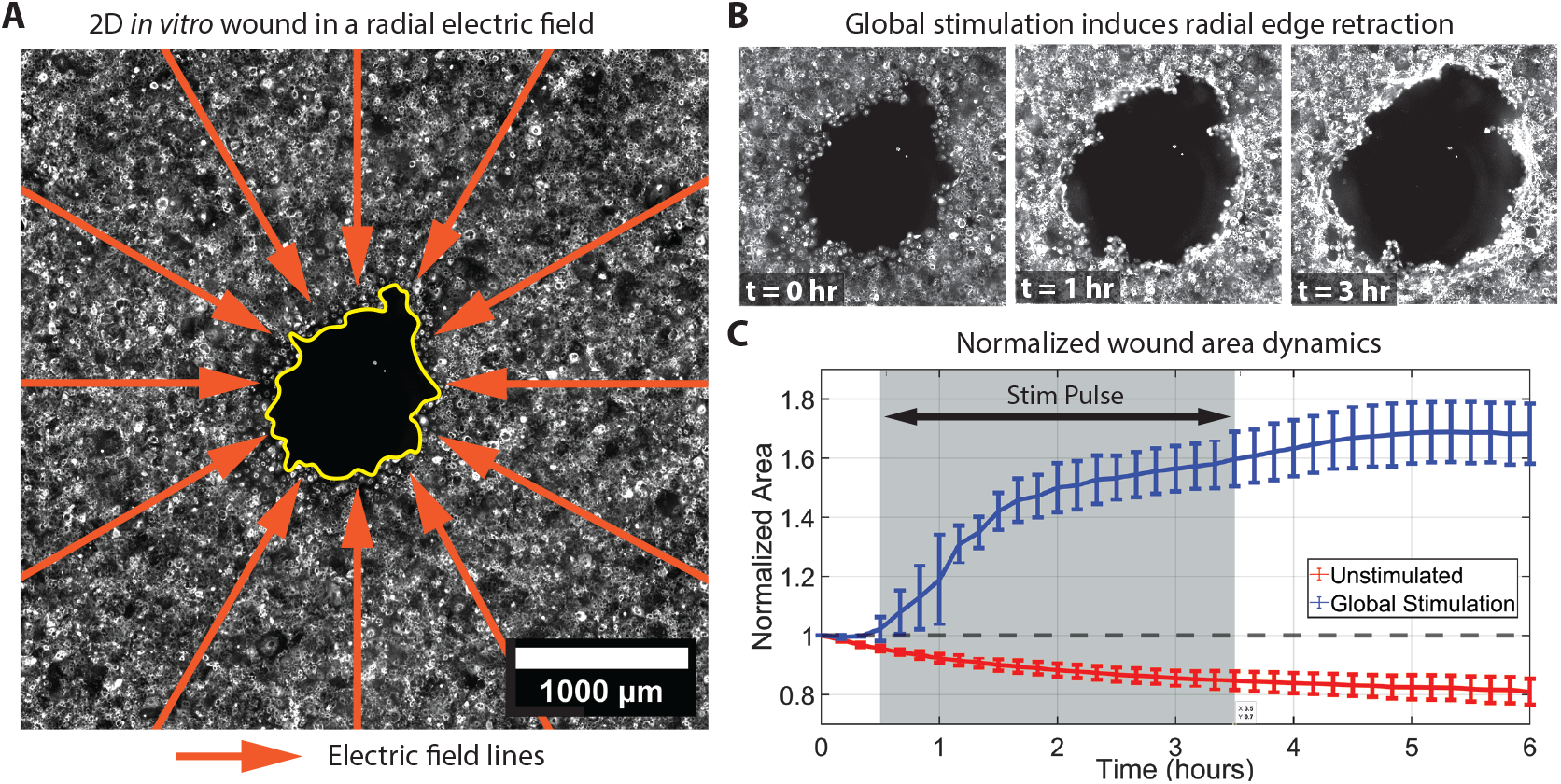
Electric stimulation induces 2D wound edge retraction. (A) Micrograph of the initial condition of a 2D *in vitro* tissue of primary mouse keratinocytes with a central wound subject to a radially inward. Electric field shown in orange and wound outlined in yellow. (B) Micrographs at *t* = 0, 1, and 3 h show edge persistent retraction of the wound boundary. (C) Normalized wound area, *A*(*t*), as a function of time compared between global stimulation and no stimulation. The dashed line at *A*(*t*) = 1 demarcates the initial normalized area of the wound and the dark grey shaded region from *t* ∈ [0.5, 3.5] h demarcates the stimulation pulse for the Global Stimulation. Unstimulated wounds heal faster because their edges do not retract.

While prior work links edge retraction to strong mechanical coupling mediated by E-cadherin [30], there is no theoretical framework to explain the failure mode. Our first goal was to develop this framework with a reductionist quasi-1D tissue geometry (1.5 mm ×6 mm strip of engineered cells), where we use an electro-bioreactor to generate a homogeneous 2.5 V*/*cm electric field instructing all cells in the tissue to move rightward [25] (see Supplemental Information section A for methods). A schematic of the device is shown in Fig. 2A and a micrograph of the tissue strip is shown in Fig. 2B. Under global homogeneous stimulation, we observe similar edge failure, as shown in Fig. 2C and in Supplemental Video 3.

**FIG. 2.**
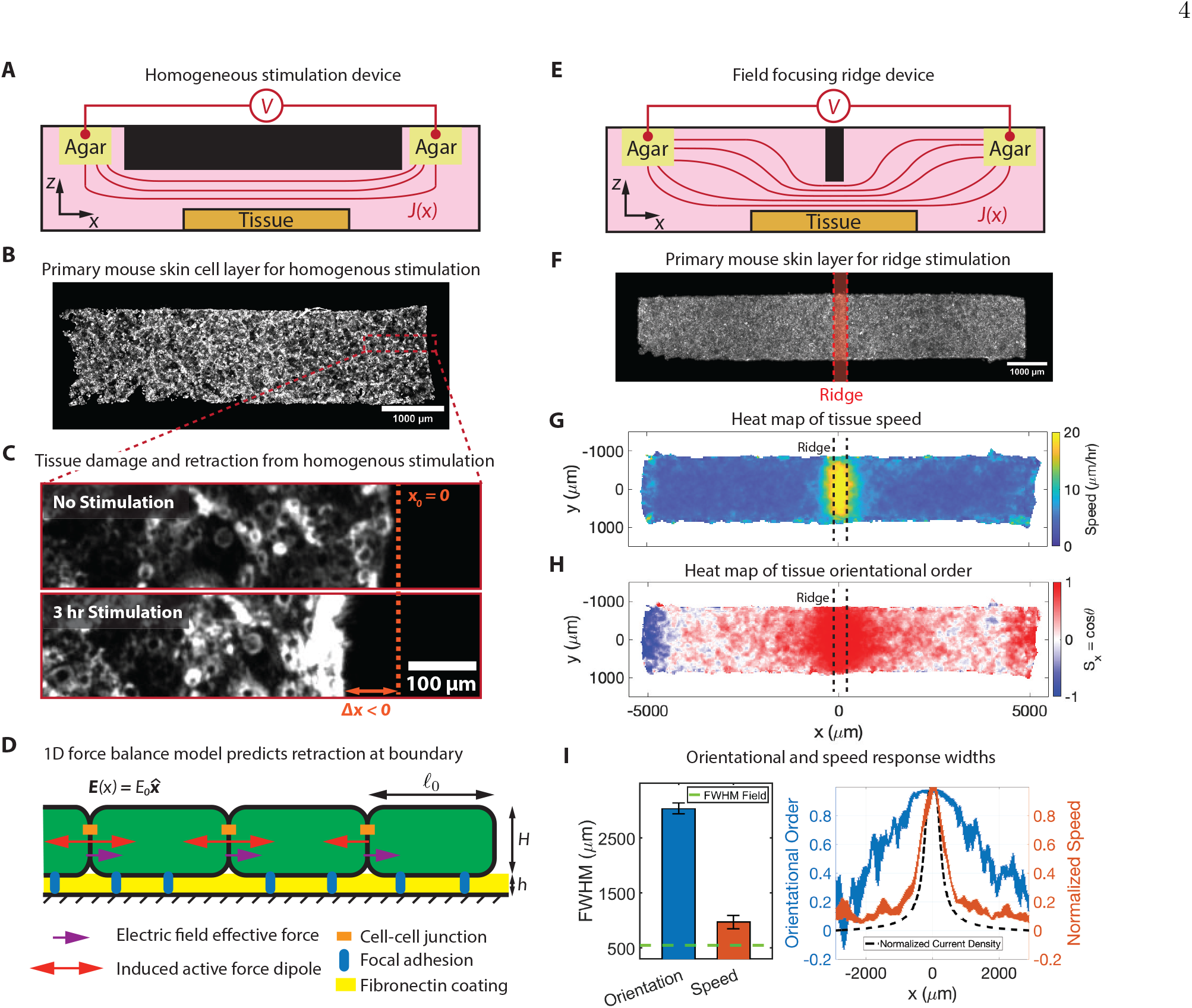
Edge retraction in quasi-1D experiments and spatially selective migration with localized electric stimulation. (A) Device schematic of 3D printed electro-bioreactor set-up for homogeneous stimulation. A voltage difference across two agarose salt bridges drives an ionic current across a tissue. (B) Micrograph of a 6.5 mm ×1.5 mm tissue strip of primary mouse keratinocytes with intermediate cell-cell coupling before stimulation. (C) Micrographs at the tissue boundary before (top) stimulation and 3 hours after (bottom) stimulation showing edge retraction. (D) 1D force balance model for a tissue in a constant electric field 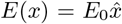. The unbalanced induced active dipole at the boundary causes retraction. (E) Device schematic of the modified electro-bioreactor used to produce inhomogeneous fields. The central ridge concentrates the ionic current above the center of the tissue. (F) Micrograph of a 10 mm ×1.5 mm strip of primary mouse keratinocytes with intermediate cell-cell coupling pre-stimulation. (G-H) Heat maps of the average (*N*≥ 6) tissue speed and the average x-component of the orientational order, *S*_*x*_, averaged over the middle hour of stimulation shows localized migratory responses. (I) (Left) Full-width-half-max (FWHM) of *S*_*x*_ and speed responses with the dashed green line indicating the FWHM of the ridge field. (Right) Unit normalized speed (orange, right axis) and orientational order (blue, left axis) averaged over the middle hour of stimulation with the unit normalized current density in dashed black.

To understand edge retraction, we build a simple continuum model shown schematically in Fig. 2D where we model the tissue as a chain of cells mechanically coupled to their neighbors and the substrate. Here, each cell has rest length *𝓁*_0_ and height *H* and is adhered to the substrate via focal adhesions through an ECM layer of height *h*. Electrotaxis acts through traditional cell migration pathways to direct motion, which we modeled as an effective force in the direction of the field, *f*_*E*_, and an active tensile force, *T*_*a*_, modeled as a force dipole. In the bulk, the net force on a cell has two spring-like contributions from its left and right neighbors, a spring-like contribution from substrate adhesion, and two effective forces from the biophysical responses to the electric field. Within the bulk, the active tensile forces are symmetric and thus cancel. Taking the continuum limit of this force balance equation results in a governing partial differential equation in the displacement 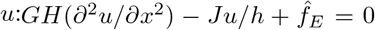, where *G* is the elastic modulus of the tissue, *J* is the substrate adhesion energy (or shear modulus), and 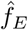 is the force density of the induced electrotactic migratory force. Unlike the bulk, at the boundary the active tensile forces are unpaired and do not cancel, leading to the boundary condition, *GH*(*∂u/∂x*)|_(*x*=*L*)_ + *σ*_*a*_ = 0, where *L* is the length of the full tissue (for N cells in the chain, *L* = *N𝓁*_0_). Provided the active stress, *σ*_*a*_ *>* 0 there is a negative strain at the edge upon stimulation, which has been shown in monolayer stress migration experiments at the edge of a tissue [24]. See Supplemental Information section C for details. This simple model demonstrates that the risk of applying global or uniform commands to a mechanically coupled tissue can arise from geometric differences between the boundary and bulk. This in turn raises the question of how to better stimulate a group and circumvent these issues. Prior work has shown that breaking cell-cell adhesion by attacking E-cadherin adhesions with drugs can improve tissue fluidity and motion [30], but this approach requires both overriding the natural mechanobiology of the tissue and weakening the integrity of cell-cell junctions which damages epithelial function and can promote infection [33, 34]. Hence, we instead focus on developing a physics-informed strategy to better apply stimuli to tissues to accelerate growth without inducing mechanical damage or requiring drugs.

### Local, spatial perturbations of a mechanically coupled group produce long-range responses

Inspired by the local perturbations in predator-prey and agricultural herding, we next investigate how applying local stimulation to only a small part of a tissue affects its larger scale mechanics. Here, we grow 10 mm ×1.5 mm quasi-1D tissue strips and redesign our stimulation platform to locally concentrate the electric field immediately over the center of the tissue spanning ~350 µm. The field strength is given by Ohm’s law, *E* = *J/σ*, where *E* is the electric field strength, *J* is the current density, and *σ* is the conductivity of the medium. To locally stimulate, we dramatically increase the current density in one region by locally reducing the height of the microchannel in which the tissue was grown (Fig. 2E and 2F). We validate stimulation structure in simulation (COMSOL) and in experiment via electrophoretic flow of charged tracer beads whose velocity scales with the current density, i.e., *J*_*x*_ ~*µv*_*x*_ where *µ* is the electrophoretic mobility. Both methods revealed a sharp Gaussian profile with a peak V*/*cm electric field (SI section A and Fig. S1 for plot) and a Full Width Half Maximum (FWHM) of 550 µm. This field is comparable to the ~1 V*/*cm endogenous field strengths measured *in vivo*.

We film cell migration for 30 minutes without stimulation (0 V*/*cm), 3 h of stimulation (2.5 V*/*cm), and 4.5 h of relaxation (0 V*/*cm) (See SI section A for methods). We quantify the induced migration patterns using particle image velocimetry (PIV) in the fluorescent membrane texture field of the cells [35] (SI section A). PIV outputs a velocity vector field, allowing us to extract the spatial profiles of cellular speed and direction as two separate metrics. Speed is computed as the magnitude of the velocity vectors (Fig. 2G), while directionality, or orientational order, is defined as *S*_*x*_ = cos *θ*, where *θ* is the angle between the applied electric field. *S*_*x*_ measures how aligned or anti-aligned the tissue motion is with the applied electrical stimuli; *S*_*x*_ = 1 (red, rightwards) corresponds to perfect alignment and *S*_*x*_ = −1 (blue, leftwards) corresponds to perfect anti-alignment (Fig. 2H).

Both the tissue speed and orientational order show a similar localized (rightward) migration of cells within the stimulation region. See Supplementary Video 4 and 5 for raw timelapses and heatmap videos, respectively. A heatmap of the divergence of the flow field shows zones of compression and extension in the vicinity of the ridge (Fig. S2 and SI section B).

This analysis reveals that locally stimulating sheets of skin with an electrotactic cue induces a hyper-local in-crease in migration speed consistent with electrotactic speed increases (Fig. 2G), but a far larger ordered domain of rightward directed migration (Fig. 2H. Red indicates right, blue indicates left). Critically, while the FWHM of the induced speed (~950 µm) is ~1.7-fold larger than that of the electrical input (Fig. 2I, red), the FWHM of the ordered domain (~3000 µm) is nearly 5.5-fold larger than the input (Fig. 2I, blue).

### Long-range migratory response is governed by continuum mechanics and cadherin-based cell-cell adhesion

What sets the length-scale over which information propagates in a mechanically coupled group? We investigate this within the framework of the previous continuum mechanics model used to describe edge retraction. In zones where the effects of electrotaxis are weak, the tissue is described by four physical parameters: the elastic modulus, *G*, the tissue height, *H*, the cell-substrate adhesion strength, *J*, and the height of the adhesive matrix layer, *h*. From these parameters, we can construct a “penetration length” scale, 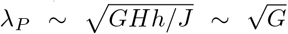, defined as the length over which information propagates via mechanical coupling beyond the Gaussian electrical input (Fig. 3A). Experimentally, we measure *λ*_*P*_ as the distance between the half-width-half-max (HWHM) of the electrical input and the HWHM of *S*_*x*_.

**FIG. 3.**
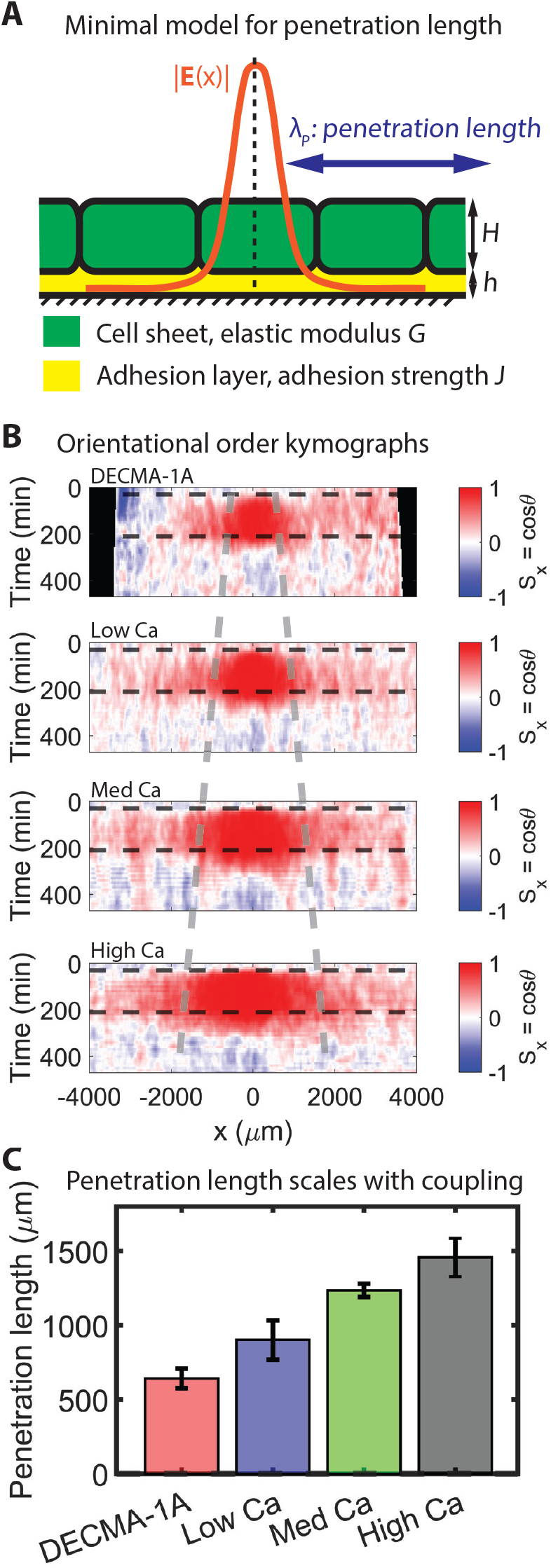
Cell-cell coupling determines penetration length scale. (A)Model set-up of tissue where a spatially localized electric stimulus produces motion that extends into the tissue bulk. The penetration length, *λ*_*P*_, is defined as the distance between the half-width-half-max (HWHM) of the stimulation pulse and that of the orientational order response. (B) Orientational order kymographs as a function of cell-cell adhesion from negligible coupling (top) to high coupling (bottom). The dashed gray lines demarcate the FWHM of the response, which grow with coupling. (C) Penetration length grows with cell-cell adhesion.

The formula for *λ*_*P*_ suggests that the penetration length scales with the elastic modulus, or effective stiffness, of the tissue, which has been shown to be strongly governed by E-cadherin mediated cell-cell adhesion in epithelial layers [36, 37]. E-cadherin proteins directly adhere neighboring cells together and act as mechanotransducers and sensors that both transmit strain and strain rate, and regulate cell stiffness by directing the cortical contractile actomyosin belt [38, 39].

We directly test this scaling model by stiffening or softening the tissue by adjusting E-cadherin levels, hoping these experiments would explain the penetration length in our data. E-cadherin adhesion strength grows with calcium levels, so we both lowered and raised calcium above the medium level (300 µM CaCl_2_) we have used for all experiments to now. As a final test, we supplement media with DECMA-1A (50 µg mL^*−*1^), an E-cadherin blocking antibody to a low calcium tissue to even further reduce E-cadherin efficacy. The results (derived from the kymographs shown in Fig. 3B) summarized in Fig. 3C show that inhibiting E-cadherin reduced the penetration length while enhancing E-cadherin raised it. For the baseline case with medium calcium levels, we measure *λ*_*P*_ ~1200 µm. In addition, we use can compute the relative moduli of tissues, *G*, which scales with cell-cell coupling as 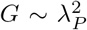 (see SI section D, Fig. S5). These tests demonstrate tissue mechanics derived from cell-cell adhesion molecular mechanisms ultimately regulate the length scale over which local inputs propagate.

### Penetration length guides 2D tissue healing and growth rates

We began by asking why inducing radially inwards migration in all cells surrounding a central wound further opened the wound. Thus far, we used quasi-1D experimental approaches and theory to explain both edge retraction and how mechanical coupling tunes the size of the collective response to a local input. We now translate these findings back to the 2D healing model, with the hypothesis that local radial stimulation might overcome the boundary retraction which occurred with global stimulation.

The simplest local, radial stimulus in 2D is an annular ring of high electric field concentric with the wound (Fig. 4A). The placement of the ring field, or the distance between the wound edge and the ring field (which we define as w), is the key experimental parameter we seek to minimize. Too close to the wound edge and the stimulus will penetrate to the edge and cause tissue retraction, but too far from the wound edge and the local stimulus will fail to affect healing. As a starting point, we use *w*~ *λ*_*P*_ ~ 1200 µm derived from the quasi-1D experiments (Fig. 3C) to guide the initial ring position.

**FIG. 4.**
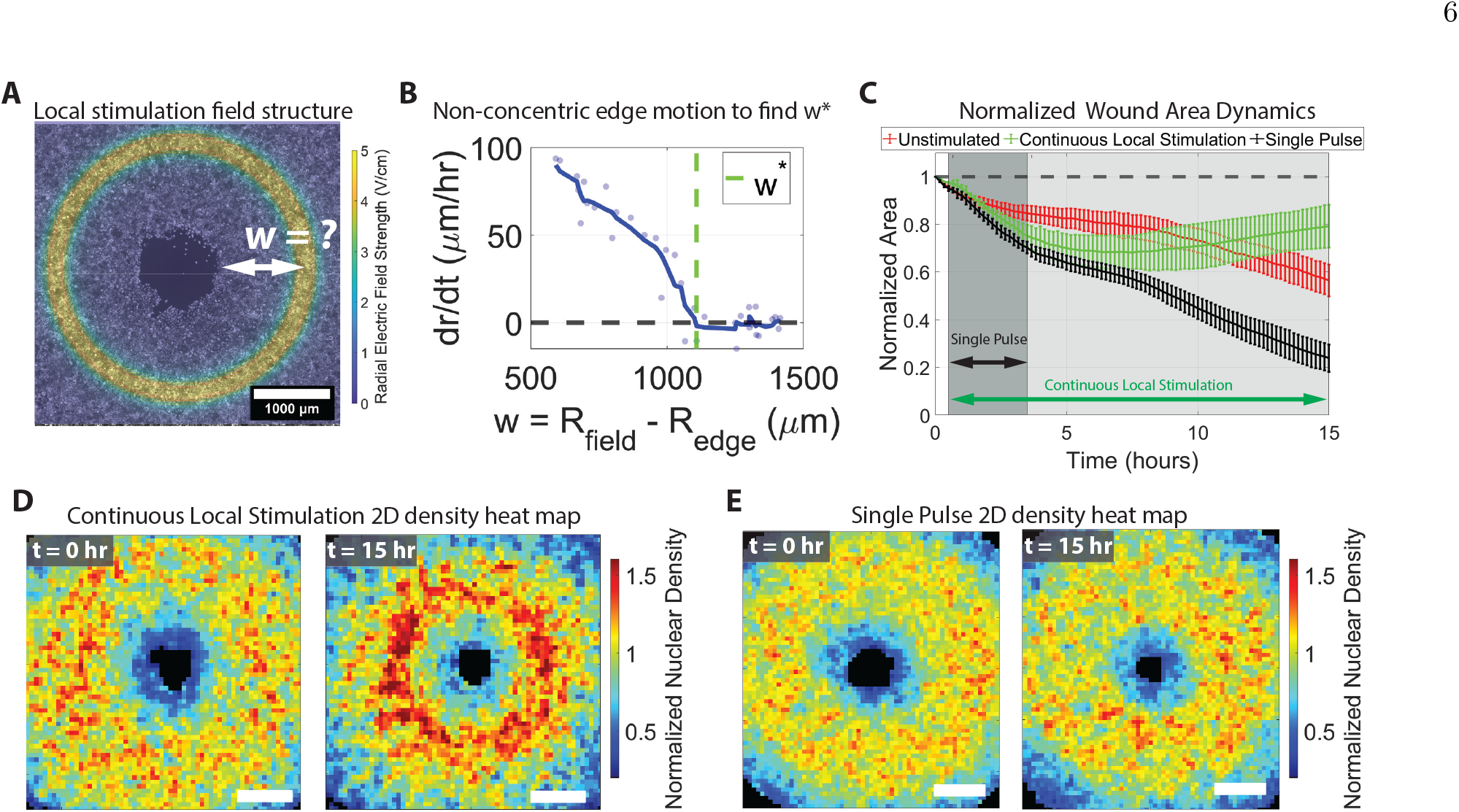
2D ring fields for wound healing are effective with pulsed fields. (A) Annular ring field for 2D wound healing assays. The distance between the edge of the wound and the field, *w*, is unknown. (B) Wound edge velocity, *dr/dt*, as a function of distance between the wound edge and the field. *dr/dt >* 0 correspond to radial edge retraction. The critical distance, *w*\*, was determined from the zero-crossing of *dr/dt*. See SI section E for details. (C) Normalized area dynamics for Continuous Local Stimulation (green), Single Pulse (black), and unstimulated (red) experiments with shaded regions and arrows denoting where the field was on for each case. (D) 2D average nuclear density heat maps from the Continuous Local Stimulation experiments at *t* = 0 and *t* = 15 h showing build-up of nuclear density from excessive stimulation. Color corresponds to local density. (F) Same as (D) but for Single Pulse data. Nuclear density never increases enough to cause jamming. Scale bars are 1000 µm.

To efficiently sweep the parameter space of possible positions, we deliberately offset the stimulus ring from the wound core. Consequently, this non-concentric ring placement generates in a single experiment a continu-ous *w*, the distance between the ring and the wound edge around the wound circumference (see SI section E and Fig. S6). Unrolling and plotting radial edge velocity, *dr/dt*, revealed a critical distance of *w*\* ~1100 µm (remarkably close to the 1D penetration length), where stimulating any closer than *w*\* would trigger edge damage (Fig 4B, zero-crossing reflects where edge retraction stops). See Supplemental Video 6 for the non-concentric stimulation assay.

### Continuous local radial stimulation triggers a jamming-like response

While continuously stimulating at *w*\* prevents edge retraction and markedly enhances early healing rates (Fig. 4C, green versus red curves), the tissue edge essentially freezes in place with longer stimulation (*>*5 h, Supplementary Video 7). We hypothesized that a simple explanation from granular active matter could explain this. To understand these dynamics, consider that the tissue is a crowd of tens of thousands of individual cells of finite size. The circular geometry and radially inward stimulation command inevitably cause cells to lock into place against their neighbors in a cellular traffic jam (an analogue to the load-supporting arch in a clogged granular hopper). To test this hypothesis, we tracked every cell in the tissue throughout the experiment with the machine-learning segmentation tool, Cellpose [40–42], and compared the spatial density profiles at the beginning of the experiment and at *t* = 15 h in Fig. 4D (see SI section F and Fig. S7). The pronounced ring of high density around the center (red), indicates where cells are likely jammed.

Knowing the slowdown emerged from a local density increase allowed us to design a solution based around the timescale of the stimulus. Tissues are active, viscoelastic materials capable of dynamically dissipating stresses given enough time, so we hypothesized that using a short duration stimulus rather than continuous stimulation might induce rapid initial migration and then allow the tissue to equilibrate and grow out before the jamming-like process occurred. The dynamics in Fig. 4C suggest that a 3 hr stimulus would have the most benefit as the plateau emerged shortly after this. We tested this new strategy, which we label “Single Pulse”, and saw that it significantly improved the healing (black curve in Fig. 4C; Supplementary Videos 8), and that it avoided density build-ups (Fig. 4E; see Fig. S7 for radial profiles). Supplementary Video 9 shows the nuclear density of the Continuous Local Stimulation and the Single Pulse stimulation together. Ultimately, the pulsed ring strategy led to 80% closure of the wound vs. 40% for the unstimulated control and 20% for the continuous ring stimulation.

### Tissue biophysics informs the optimal stimulation strategy

Our short pulse, long-term relaxation strategy significantly improved tissue healing, but if one pulse is good, would more pulses be better, and if so, where should we apply them since the wound is shrinking dynamically? We first test this by applying a second pulse in the same position as the first with a 6.5 hour relaxation period between pulses, but found this did not improve healing (Supplementary Video 10). We reasoned that this is related to how the wound shape and the tissue mechanics are changing over time, and that a better solution would be to adapt the location of the stimulus to the current shape and size of the wound.

Given the large parameter space of possible pulse sequences and locations, we built a continuum model based on physics of an active tissue embedded in an optimal control framework (full details in the SI section G) to help refine our control strategy. This optimal control instantiation, based on the Hamilton-Jacobi equation, has deep connections to classical physics. Briefly, the physics of the experimental system is described by a density of active cells that vary in space-time, *ρ*(*x, t*), and an accompanying velocity field, *v*(*x, t*), which is subject to control by an external electric field, *u*(*x, t*). Here we assume the dynamics can be described in 1 space dimension (or in an axisymmetric geometry) for simplicity. Assuming that mass is conserved over the course of an experiment, which we verify using experimental measurements of cell counts, the continuity equation reads

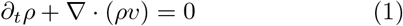

In a minimal setting of wound healing of an epithelial gash, cells respond to variations in density by moving to close the wound but maintain their integrity. Their motion is resisted by friction with the substrate and with each other, and driven by a combination of density variations and the external electric field, leading to a minimal equation for force balance that reads

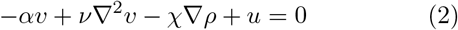

Here *αv* represents friction with the substrate, *v* ∇^2^*v* represents viscous forces due to mechanical coupling within the sheet, −*χ* ∇*ρ* drives motion due to density gradients with a demotactic (Etym., *demos* = population, *taxis* = directed movement) coefficient *χ*, and *u*(*x, t*) is the local control induced by the applied electric stimulus that is as yet unknown.

To determine this control field, we postulate the control objective as one that minimizes the wound area, i.e., uniformizes the cell density field when starting from a wounded state, subject to an *L*^1^ norm cost of the controller, that physically corresponds to a sparseness cost in space-time (see SI section G), and corresponds to the following formulation of the optimal control problem:

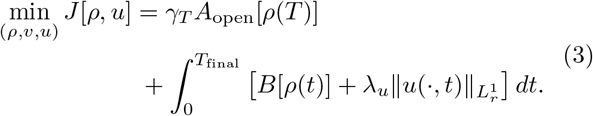

where the cost function, 𝒥 is composed of a terminal cost (*γ*_*T*_ *A*_open_[*ρ*(*T*)]) which penalizes the wound area and a running cost 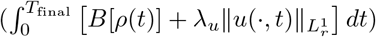 which penalizes running the controller and is minimized subject to the mass and force balance PDE constraints. The normalized epithelial density is modeled by a sharp tanh(*x*) function at the wound boundary.

We solve the stochastic optimal control using a recently introduced adjoint-based approach to solve the Hamilton-Jacobi-Bellman equation, that uses a path integral formulation and continuous-time back-propagation to evaluate the control strategy in terms of the value function [43–47] (see SI section G). The solution predicts that the most effective control strategy for accelerated healing is to apply a series of pulses which dynamically move along with the wound edge as it heals. This result, shown in Fig. 5A as a kymograph, demonstrating that as the wound dynamically shrinks, so must the ring stimulus.

**FIG. 5.**
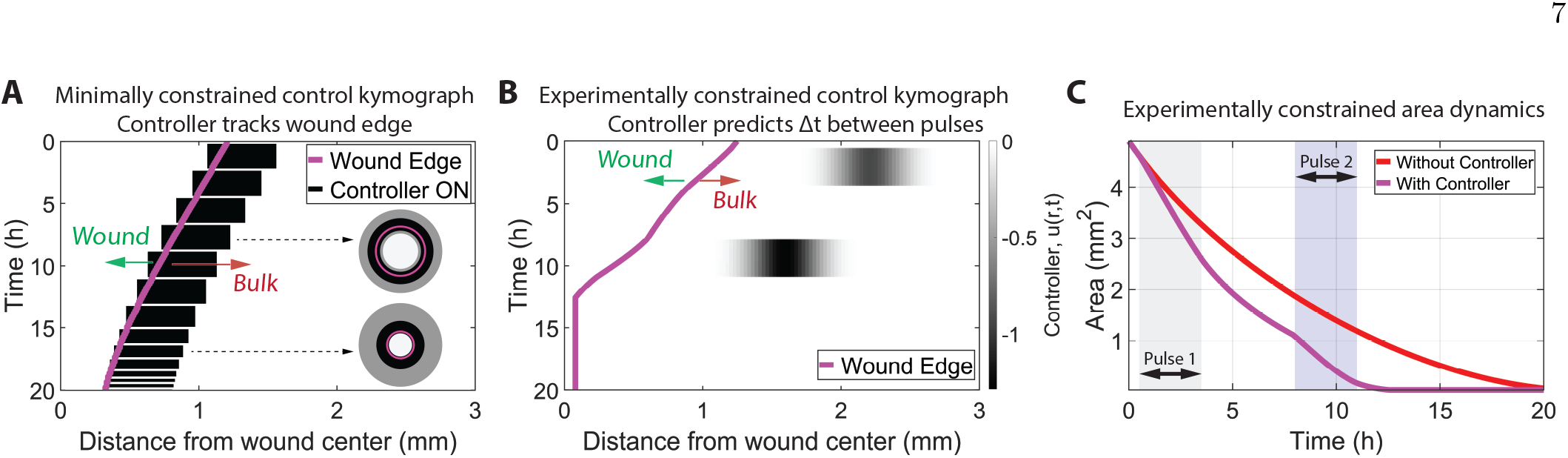
Optimal control solution in minimally constrained and experimentally constrained limits. (A) Kymograph of minimally constrained optimal control policy predicts a pulse train that tracks the wound edge in space as it heals in time. The pulses have fixed magnitude, *u*(*r, t*) = −0.1. The wound edge is plotted in purple with arrows indicating the wounded regions (*ρ* = 0) and bulk regions of finite mass (*ρ >* 0). Inset: snapshot schematics in time of two ring pulses which track the wound boundary. (B)Experimentally constrained optimal control solution yields a double pulse sequence with pulses offset in space and time. The wound boundary is plotted in purple. (C) Simulated wound area for the experimentally constrained optimal control double pulse sequence (purple) and the naturally healing case (red) without a controller.

While the optimum strategy is a nearly continuously spatially varying stimuli that tracks the healing edge, testing this physical prediction experimentally requires additional constraints. First, the human experimenter needs to manually change the ring stimulus by replacing the initial ring stimulator with one of a smaller diameter. This experimental challenge meant we could only practically evaluate two ring diameters. Next, we needed to encode an explicit penalty in the cost functional *B*[*ρ*(*t*)] based on our experimental knowledge of the system. We thus penalized bringing the controller closer than a dis-tance of *w*\* to the wound edge which causes wounds to open further. Finally, we restrict the simulation to 20 hours, a few hours longer than the experimental time to minimize the role of cell division in the process. The cell cycle for the mouse keratinocytes is about 16 hours, and we verified that there were no significant changes in cell counts over this duration.

With these constraints in place, the optimal control problem yields a solution for where to place the two annuli in space and how long to wait between stimulation pulses (see SI section H for details). In Fig. 5B, we plot control kymograph to the experimentally constrained optimal control problem, which predicts two, 3 h pulses at *R*_1_ ≃ 1.1 mm and *R*_2_ ≃ 0.75 mm separated in time by Δ*t* = 6 h where the second pulse is stronger in magnitude. Fig. 5C shows the area of the electrically stimulated wound compared to the case in the absence of an electric field, demonstrating the speed-up of 40% for wound-closure. Accompanying movies of the wound density and controller dynamics for the experimentally constrained case and the case without control are shown in Supplementary Video 11.

### Experimental Double-Pulse strategy improves healing 5-fold

We take inspiration from these theoretical predictions and test them experimentally using our final control strategy that we call “Double Pulse”. To clearly demonstrate this schema, we show a schematic set-up in Fig. 6A, where two annular ridges of radii *R*_1_ = 1.75 mm and *R*_2_ = 1.5 mm representing the two pulses overlay a tissue micrograph. Between the two pulses there is a time delay of Δ*t* = 7 h. A kymograph for this control strategy and a video of the Double Pulse experiment are found in in Supplementary Fig. S10 and Supplementary Video 12, receptively.

**FIG. 6.**
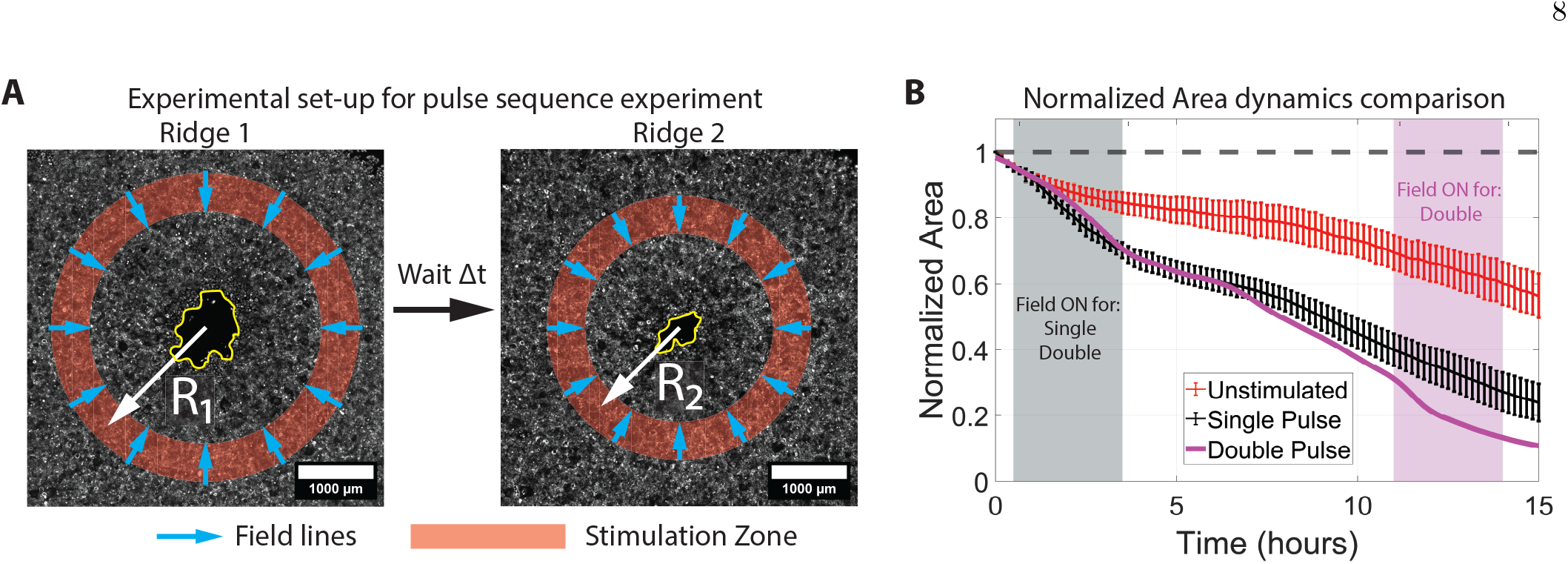
Experimental realization of constrained optimal control solution shows faster wound healing. (A) Schematic of the double pulse sequence inspired by the predictions of the constrained optimal controller. Two annular ridge fields with radii *R*_1_ and *R*_2_ are overlaid on wounded tissues with wound boundaries in yellow. (B) Normalized area dynamics of the double pulse data compared to the single pulse data with shaded regions for the stimulation pulse periods. After pulse 1, the two traces track one another well. However, pulse 2 accelerates the healing rate again which splits the two traces. The unstimulated strategy is plotted in red for comparison.

As before, we evaluate the controller performance by plotting the healing dynamics in Fig. 6B. We include the results from the Single Pulse experiments as a point of comparison and the unstimulated wound as the baseline. We see two regions (corresponding to the two pulses) where healing is accelerated via the optimal control generated stimulation pulses. The Double Pulse strategy far outperforms all other strategies, and it represents a nearly 5-fold increase in healing rates over the Unstimulated strategy. Videos of various alternative instantiations of Double Pulse sequences (Supplementary Videos 10 and 13) both underperform the optimal solution.

## DISCUSSION

We began with the broad question of how best to leverage crowd-informed physics to control collective motion, using the example of electric-field accelerated healing a model skin wound as a case study. These skin tissues were an ideal model system since they have tunable collective coupling through cell-cell adhesion and guidable motion through electrotaxis. Our results, enabled by a newly designed local bioelectric stimulation tool, revealed certain powerful guiding principles derived directly from the biophysics of crowds. In particular, where and when stimuli are applied can radically influence crowd mechanics and migratory dynamics. These concepts demonstrate that control strategies are limited by how well we can understand the physics behind the crowd, but building the physics into a controller can allow us to synergize with the natural mechanics to achieve a goal rather than needing to override or break the natural mechanics [30].

We first found that where a stimulus is applied (global versus local) radically affects the resulting mechanical response. Global inward stimulation of cells in a tissue resulted in edge damage attributable to differences in the boundary and bulk of a tissue. We were able to circumvent tissue damage with a highly localized stimulus, inspired by how a sheepdog controls a flock of sheep by applying local pressure that propagates from the outside of the flock inwards. Our thin electrical pulse applied within the bulk mechanically propagates to the boundary, from inside out, over a long length scale set by cell-cell adhesion (the penetration length derived in Fig. 2). However, we also found that local perturbations are insufficient for effective group control of tissue mechanics and when the stimulus is applied matters greatly. In particular, a continuous local inward stimulus gradually induced a massive increase in local density, reminiscent of a jamming transition but likely more complex, that ultimately quenched healing. Simply allowing the tissue to dissipate stress and relax, taking advantage of its effective viscoelasticity, bypassed jamming. This problem also highlights another key detail of crowd mechanics—the boundary conditions and geometry dramatically affect local control. Here, the circular geometry exacerbated the jamming-like behavior because cells converged radially to a point. In uniaxial motion, jamming is considerably less likely.

Finally, we demonstrated that where and when stimuli are applied is a dynamic decision that relies on understanding the biophysics, geometry, and boundary conditions of the system. Delivering two local pulses of stimulus at the same spatial location separated by a relaxation period does not improve tissue healing. By contrast, delivering two local pulses at different positions chosen to track the advancing wound edge was much more effective than a single pulse of stimulation. Reaching this conclusion required developing a physics-based control model which accounted for practical experimental limitations then optimized where and when to pulse the field. This physics-derived optimization strategy is particularly compelling as it demonstrates how a minimal physical model of collective dynamics can be coupled to optimization strategies that ultimately allow precise external control of collective mechanics.

While we articulated these basic principles of crowd regulation, there are many limitations we wish to highlight that could lead to exciting future work. First, we focused on a skin model, and how these strategies work in different systems will depend heavily on how agents within the collective couple, which varies by system. Second, our data reveals rich physics that we cannot fully explain. For instance, there is a local mechanical recoil when local stimulation stops (Fig. 3B). It is unclear where this comes from as the timescale is too long for elastic recovery. Similarly, the gradual, freezing of the leading edge of healing tissues during continuous local stimulation is also surprising and likely reflects an additional hidden length scale in the group dynamics.

Our findings may have implications for future work across different crowd systems. One example is accelerating actual injury healing *in vivo*. A growing collection of studies all point to beneficial effects of trying to apply directional electric fields to injuries in animal models [48–50], but no standard strategies for where and when to stimulate have been articulated and all systems to date apply global, instantaneous fields. It is possible that future work adapting our spatial and temporal stimulation strategies here could help to optimize performance in these applications. Beyond tissue systems, the rules we demonstrate here may generalize to other crowds, such as dense human crowds where dynamic, local signage and guidance may help to improve foot traffic flow or reduce the chance of mass crowd crushes where global commands can cause more confusion than good.

## ACKNOWLEDGMENTS

We thank The Eric and Wendy Schmidt Transformative Technology Fund, National Institutes of Health Award R35 GM133574-07, and National Science Foundation CAREER Award 2412942 (DJC), The Omenn-Darling postdoctoral fellowship (JSY), The Simons Foundation and Henri Seydoux Fund (LM) DARPA MINT (SS, VK, LM). JSY thanks S. Ganga Prasath for useful conversations.

## AUTHOR CONTRIBUTIONS

J.S.Y. and Y.L. performed experiments. J.S.Y. analyzed results. S.S. and V.K. performed optimal control modeling. J.S.Y., L.M., D.J.C. conceived of project and analysis. J.S.Y., L.M., and D.J.C. wrote manuscript with comments from all other authors.

## DATA AVAILABILITY

Key raw microscopy data and figure-generating data from the main text will be available on Zenodo (https://zenodo.org/records/17673840). The code for the optimal control analysis is accessible through GitHub (https://github.com/sumit-sinha-seas/wound_healing_optimal_control.git).

