## Supplementary Information for "Locally stimulating cell migration in living tissues drives long range collective motion through cell-cell adhesion leading to accelerated migration, healing, and growth"

**D Effective elastis modulus of tissues as a function of cell-cell coupling** **9**

**E Non-concentric wound healing assay for finding edge retraction length scale** **9**

**F Normalized nuclear density line profiles, healing rates for different stimulation conditions, double pulse kymograph** **10**

**G Optimal control for emergence of pulsed strategy: minimally constrained** **13**

**H Experimentally constrained optimal control: Brief problem description** **16**

**I Description of Movies** **21**

### A Methods:

#### A.1 Cell maintenance

Primary mouse keratinocytes (a gift from Prof. D. Davenport, Princeton University) were cultured in E-medium (Nowak and Fuchs, 2009) supplemented with 15% fetal bovine serum. To tune the coupling between cells, $CaCl_2$ was added to the growth media at a concentration of 50 µM (low), 300 µM (medium) or 1000 µM (high). In addition, coupling could be damaged via by adding DECMA-1A, an E-cahderin targeting antibody. Cells were maintained at 37° C with 5% $CO_2$ and 95% relative humidity. Cells passage numbers were kept below 30 for all experiments.

#### A.2 Cell stenciling and substrate preparation

Tissues were seeded into standard plastic 24-well plates (EW-01927-74, Cole-Parmer) where each well was coated with 95 µL of 50 µg mL$^{-1}$ fibronectin (FC-010, Millipore Sigma) under a 15 mm circular glass coverslip (CLS-1760-015, Chemglass) for 30 min at 37 °C, then washed three times with deionized water (DI). Next, polydimethylsiloxane sheets approximately 250 µm thick (Bisco HT-6240, Stockwell Elastomers) were cut using a stencil cutter (Silhouette) into various shapes ranging from a strips to disks and placed into the stimulation wells (typically columns 2 and 5 of the 24-well plate; D4, D5, and D6 served as negative control wells).

Cells were counted using an automated cell counter (Cytosmart, Corning) and seeded at a density of $2.25 \times 10^6$ cells/mL then a micro volume, $V$, computed via $V = A \times 0.41$, where $A$ is the open stencil area is seeded into the micro-stencil. The circular border of each target well was lined with approximately 100 µL of media to mitigate evaporation. After approximately 4 h in the incubator, once cells had fully attached to the substrate, 500 µL of media was supplemented to each well. Stencils were removed approximately 16 h after incubation. Timelapses began 1 h to 2 h after stencils were removed.

#### A.3 Circular wound assay stenciling protocol

We first placed a 1.5 mm diameter circular PDMS disc in the center of each stimulation well, followed by a second circular PDMS stencil with a 7 mm diameter circular cut-out concentric with the 1.5 mm inner disc. This created a large circular reservoir with a small disc in the center.

We next seeded cells in low-calcium (50 µM $CaCl_2$) media. After cells attached to the substrate (approximately 4 h to 6 h), we rinsed twice with PBS and pulled off the central stencil using tweezers. This mitigated any cells in suspension from landing and adhering to the wound region. We then rinsed once with medium-calcium (300 µM $CaCl_2$) media, and subsequently flooded the wells with medium-calcium media for approximately 24 h. Importantly, the void in the center began to fill in overnight through proliferation and migration, yielding wounds typically between 0.5 mm to 1 mm in diameter ($0.5 < d < 1$ mm). Although wounds initially formed circularly, by the start of experiments they often exhibited a more heterogeneous edge profile.

##### A.4 Imaging Preparation and Microscopy Set-up

Cells were stained with a cytoplasmic membrane dye, CellBrite Red (Excitation / Emission 644 nm / 665 nm; 30023, Biotium) diluted 5:1000 in media, and the nuclear stain Hoechst 33342 diluted 1:2000 in media, then incubated for 30 min at 37 °C and triple-rinsed in media. A BSA fiducial marker in the Cy5 channel (B5S, ProteinMods) used for registration was placed in the vicinity of each tissue. Images were acquired with an inverted fluorescent microscope (Ti-2, Nikon) equipped with a cage incubator that maintains 37 °C and 5 % humidified $CO_2$ using a $CO_2$ bubbler. Automated motorized stage and capture systems were controlled using Nikon Elements (Nikon), and a CMOS camera (Qi-2, Nikon) was used for image acquisition. Most experiments used a 4× 0.2 NA objective.

Images were captured in the Cy5/DAPI channels with typical imaging parameters of 10 min capture frequency at 15/10 % lamp power (Sola SE, Lumencor, USA) and 400 ms exposure. For multi-point images, a 10 % image overlap was used. For radial wound healing assays (day-long timelapses), we used gentler imaging conditions were used: the Cy5 channel was captured with 10 % laser power and 200 ms exposure every 10 min, and the DAPI channel was captured with 5 % laser power and 400 ms exposure every 30 min.

##### A.5 Electrotaxis Experimental Set-up

The electro-bioreactor set-up follows [1]. Each stimulation insert was 3D printed using white PLA filament (Polylite PLA natural, Polymaker) on a 3D printer equipped with a 0.2 mm nozzle (X1C, Bambu Lab) and slicer settings of 90 % infill, 0.08 mm layer height, with supports enabled. The high infill ensures that inserts sink into each well. The supports were manually removed, and each insert was dialyzed in approximately 1 L of sterile deionized (DI) water overnight.Agar salt bridges were prepared by melting agarose (20-102, Apex Bioresearch) and PBS (D8537, Sigma-Aldrich) at 4 % weight/volume on a hot plate at 180 °C, after which the molten agar was cast acrylic molds. Ag/AgCl electrodes were prepared by plating thin strips ∼8 mm × 23 mm of silver foil (AA11440GW, Fisher Scientific) in 1 M KCl solution (P9541-1KG, Millipore Sigma) for 18 h at 1 mA using a titanium wire cathode (00362-G1, Alfa Aesar). Note that Ag/AgCl is a non-polarizable electrode, so under high voltages, it generates redox products. The salt bridges in the assembly act as an electrochemical barrier which protects the sample.

Inserts, electrodes, and agar bridges were sterilized under UV exposure for 5 min in a biosafety hood. Wells adjacent to the tissue well were filled with ∼2 mL of PBS each, and approximately 1.5 mL of media was added to each tissue well. Inserts were then placed into the corresponding wells, and agar bridges were positioned on top of the inserts. Electrodes were inserted into the PBS wells. Finally, an incubator lid was mounted on top of the assembly for live imaging.

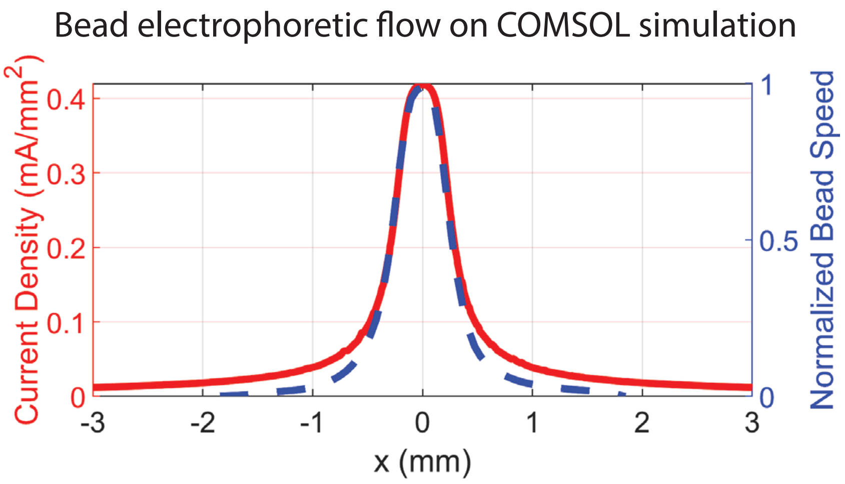

Figure S1: COMSOL simulation of the current density in the x-direction (red) showing strongly localized current density. Overlaid (blue dashed line) is the normalized x-component of the velocity of charged beads flowing in ridge device showing that the COMSOL simulation accurately scales with the ground truth.

#### A.6 Bead Validation and COMSOL Simulations

Finite element analysis simulations (COMSOL 5.6) were used to generate preliminary predictions for the 1D ridge stimulation current density using a material conductivity of $1.6\,\mathrm{S\,m^{-1}}$ and relative permittivity of 80. To experimentally measure the current density, two Ag/AgCl-tipped Ag wire probes were held at bottom of the insert, and the device cross section was measured using confocal microscopy (NL5, Confocal NL). The read-out voltage agreed well with the COMSOL simulation. Temperature increases from Joule heating were measured by placing a T-type thermocouple (5TC-KK-T-30-72, Omega) at the bottom of the well while running stimulation. The temperature readout (54 II B, Fluke) changed by less than 1 °C, indicating Joule heating was negligible.

To validate the non-uniform ridge field geometry, 1 µm diameter fluorescent charged beads (F-8765, Thermo Fisher) were diluted 1:1000 in PBS then electrophoretically driven across the device. The setup was imaged using confocal microscopy with a 10× 0.3 NA objective in the FITC channel at 5 frames per second. The velocity vector field of the charged beads was computed with particle image velocimetry (PIV) analysis (described in Section A.7.2), with care taken to subtract background flows. At intermediate field strengths, the speed of electrophoretic flow scales with the drag force on the charged bead in a viscous, conducting, medium scales, i.e., $v = \mu E$ where $\mu$ is the electrophoretic mobility. Thus, the experiment generates a map connecting field strength to velocity, $v \sim E$ (up to scaling). Fig. S1 shows the normalized $v_x$ overlayed on the current density results from the COMSOL simulation, and indicates an effective electrical stimulation zone of full width half max (FWHM) $\sim 550\,\mu\mathrm{m}$ mm with a peak current density of $J_x = 0.42\,\mathrm{mA\,mm^{-2}}$ (corresponding peak electric field strength of $E \sim 2.5$ V/cm)

#### A.7 Microscopy Imaging and Analysis

##### A.7.1 Pre-processing in Nikon Elements and ImageJ

All microscopy images were stitched using either the built-in stitching algorithm in Nikon Elements or the Grid/Collection stitching algorithm in ImageJ, with a 10% tile overlap and linear blending

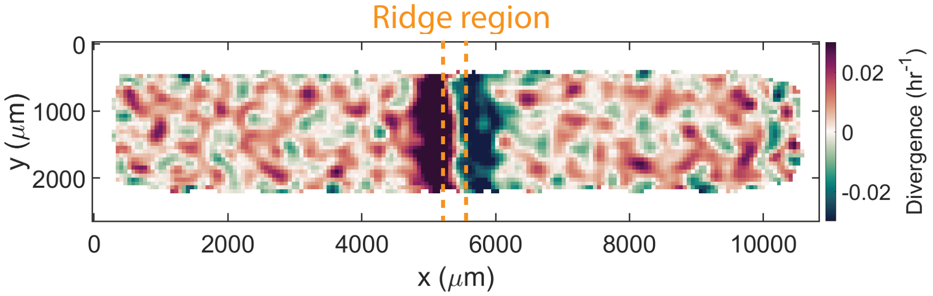

Figure S2: Heatmap of the average divergence of the velocity field taken from the middle hour of stimulation which shows the localized migratory response of the tissue averaged over $N \geq 6$ samples.

at the boundaries. To eliminate drift, images were registered across time points to the fiducial BSA marker in the Cy5 channel using ImageJ's built-in template matching algorithm with subpixel accuracy. Finally, image backgrounds were masked in ImageJ using a combination of Gaussian and median filtering, thresholding, and moving-average background subtraction.

##### A.7.2 Particle Image Velocimetry (PIV) Set-up

The velocity vector field of cell migration motion was obtained using the MATLAB plugin PIVlab [2] using three passes, each with a 50 % step overlap: first a 128 × 128 pixel interrogation window, then a 64 × 64 pixel window, and finally a 32 × 32 pixel window. Outliers, defined as those exceeding 8 standard deviations from the mean, were filtered and replaced with interpolated values.

The velocity vector fields are exported component-wise ($u_x$ and $v_y$) then imported into MATLAB and used to compute speed and orientational order. The speed, $|v|$, was computed by vectorially summing $u_x$ and $v_y$ at each voxel and then computing the magnitude, i.e., $|v| = \sqrt{u_x^2 + v_y^2}$. The orientational order was computed via $S_x = \cos\theta = u_x/|v|$. This parameter measures how aligned the migratory motion is with the electric field (here, $E \sim E_0\hat{x}$). Note, in the maintext, the orientational order is sometimes referred to as "directionality". These scalar fields were the basis for generating heat maps and kymographs. Kymographs were computed by first averaging vectors through the bulk of the tissue along the vertical direction, avoiding the upper and lower edges, then stacking each 1-D array according by time.

##### A.7.3 Cell Segmentation and Nuclear Density Computations

To calculate nuclear density in the radial wound healing experiments, we segmented nuclei using Cellpose [3]. Although the built-in model, CPSAM, performs well on our datasets, we found that training a custom model yields higher accuracy. Typically, we select 10–20 reference images (covering a small portion of the larger tissue in the DAPI channel), manually label them, and train a new model initialized from CPSAM. The model is then used to batch-process the entire image stack, outputting the segmented coordinates of each nucleus. Our results were also sensitive to certain basic Cellpose parameters. The specific settings we used were within the following ranges: diameter $\in [10, 20]$, flow threshold $\in [2, 3]$, and cell probability threshold $\in [-6, -4]$. We validated the segmentation by overlaying the Cellpose segmented centroids of nuclei on the ground truth nuclear

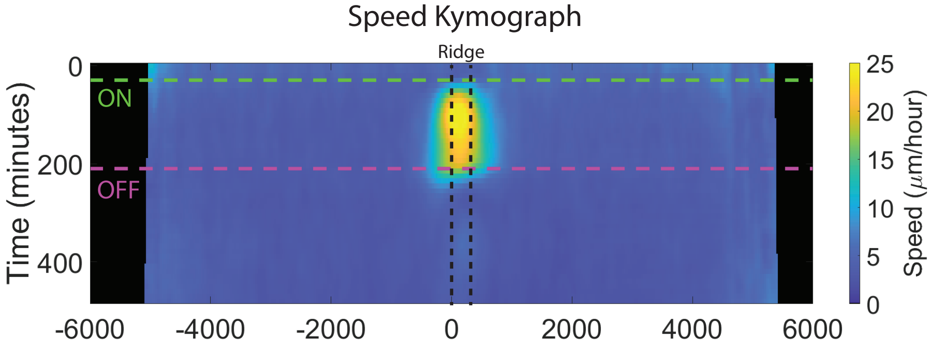

Figure S3: Kymograph of the average speed from N = 6 medium Calcium keratinocyte tissues with color corresponding to speed. The green and purple dashed lines indicate when stimulation is turned on (30 minutes) and off (210 minutes) respectively.

channel images for each data set.

#### B Divergence heat map, speed kymograph, and transfer function for ridge field

For the quasi-1D experiments, we use the vector fields derived from PIV to compute the divergence of the flow. In Fig. S2, we see regions of compression and expansion in the ridge region separated by a seam where flows match and the divergence vanishes. Since the velocity is small in the $y$-direction, the divergence and strain rate are nearly identical.

Kymographs of orientational order can be found in the maintext. Here we include a kymograph of the speed for Medium cell-cell coupling computed directly from PIV data by projecting speeds onto the $x$-coordinate taking care to avoid the upper and lower boundaries of the tissue are plotted in Fig. S3 (averaged over $N \geq 6$ tissues). Dashed lines indicate where stimulation was turned on and off. We observe that after stimulation begins, the migration speeds quickly (within $\sim 30$ minutes) broaden to their final spatial extent of with a FWHM of 950 µm.

From the speed and orientational order kymographs, we compute an input-output response, or transfer function, which describes how the tissue speed and orientational order map to current density (in our experiments, the current density is continuous in the $x$-direction). Please note that the transfer functions are specific to this experimental setup and are functions of field geometry, tissue type, and cell-cell coupling. They should not be interpreted as maps which would apply to homogeneous stimulation, for instance. The transfer functions for orientational order and tissue speed (raw values) are plotted in Fig. S4. As the current density increases, the tissue speed gradually and monotonically increases from its baseline to its peak speed. The speed response is weakly nonlinear in current density within this domain. In contrast, the orientational order, rapidly increases and saturates to its maximum at relatively low current densities ($J_x \simeq 0.1$ mA/mm$^2$). This saturation is a consequence of mechanical coupling.

#### C Derivation of the continuum model

The simplest method to derive the continuum equations is with a discrete spring model of cells. Consider a 1D chain of $N$ cells with average rest length $\ell_0$; the total length of the chain is thus

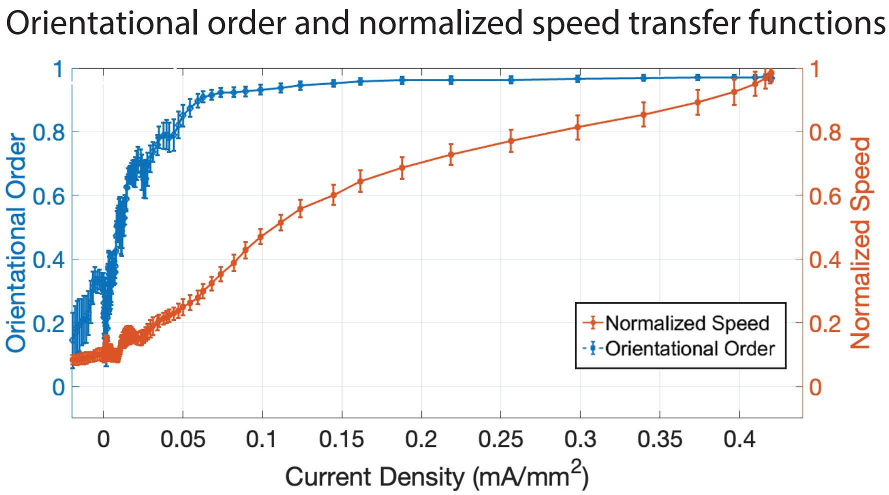

Figure S4: Normalized speed (orange, right axis) and orientational order (blue, left axis) as a function of input current density. Data is the average response from the middle hour of experiment and is derived from the kymographs found in Fig. S3.

$L = N\ell_0$. The resting position of the $i^{\text{th}}$ cell is $x_i = i\ell_0$. Under deformation, the actual position of the $i^{\text{th}}$ cell becomes $x_i + u_i$, where $u_i$ is the local displacement. We model the mechanical forces using the following four components:

1. Cell-Cell elasticity: Cells are connected to their neighbros by springs with spring constant $k_1$, which to first order describes the amount of E-Caherin in the cell.
2. Substrate adhesion: Each cell is anchored to the underlying extra-cellular-matrix (ECM) by a spring of stiffness $k_2$.
3. Active dipole-like force: The electric field induces an internal actomyosin contraction. We model this as an active force dipole wherein the electric field shortens the bond between cell $i$ and cell $i+1$ generating an active tension $T_a$.
4. Electrotactic mobility: The electric field induces a phenomenological force, $f_E$, along the field.

#### C.1 Description in tissue bulk

The net force on the $i^{\text{th}}$ cell in the bulk of the tissue has spring contributions from two neighbors, the substrate spring, and two forces induced by the electric field,

$$F_i = [k_1(u_{i+1} - u_i) + T_a] - [k(u_i - u_{i-1}) + T_a] - k_2 u_i + f_E = 0 \tag{1}$$

Provided $T_a =$ constant these cancel, so,

$$F_i = k_1(u_{i+1} - 2u_i + u_{i-1}) - k_2 u_i + f_E = 0 \tag{2}$$

In the continuum limit,

$$u_{i+1} \approx u(x) + \ell_0 \frac{\partial u}{\partial x} + \frac{\ell_0^2}{2} \frac{\partial^2 u}{\partial x^2} + ... \tag{3}$$

$$u_{i-1} \approx u(x) - \ell_0 \frac{\partial u}{\partial x} + \frac{\ell_0^2}{2} \frac{\partial^2 u}{\partial x^2} - ... \tag{4}$$

Therefore,

$$u_{i+1} - 2u_i + u_{i-1} \approx \ell_0^2 \frac{\partial^2 u}{\partial x^2} \tag{5}$$

Substituting into Equation 2 gives the force at a position $x$ within the bulk,

$$F(x) = k_1 \ell_0^2 \frac{\partial^2 u}{\partial x^2} - k_2 u + f_E = 0 \tag{6}$$

To convert this to a continuous force density (force per unit length), we divide the entire equation by the cell length $L$:

$$(k_1 \ell_0) \frac{\partial^2 u}{\partial x^2} - \left( \frac{k_2}{\ell_0} \right) u + \left( \frac{f_E}{\ell_0} \right) = 0 \tag{7}$$

Mapping to continuum quantities: $k_1 \ell_0 \to GH$ where $G$ is the elastic modulus, $H$ is the height of the cell, $k_2/\ell_0 \to J/h$ is the substrate friction where $J$ is an adhesion energy or shear modulus and $h$ is the height of the ECM layer/adhesive layer, and $f_E/\ell_0 \to \hat{f}_E$ is a continuum force density from electrotaxis. Therefore,

$$GH \frac{\partial^2 u}{\partial x^2} - \frac{J}{h} u + \hat{f}_{E} = 0 \tag{8}$$

#### C.2 Description at the leading boundary

At the boundary of the tissue (the $N^{\text{th}}$ cell), there is a half-dipole to the left connecting to cell $N-1$, but not to the right. The force is now unbalanced on cell $N$,

$$F_N = 0 - T_{N-1,N} - k_2 u_N + f_E = 0 \tag{9}$$

$$- \left[ k_1 (u_N - u_{N-1}) + T_a \right] - k_2 u_N + f_E = 0 \tag{10}$$

Assuming the boundary layer is infinitesimally thin, the substrate drag and body forces scale with volume and vanish at the boundary. The equation strictly reduces to a balance of boundary tensions,

$$k_1 (u_N - u_{N-1}) + T_a = 0 \tag{11}$$

Expanding to first order $(u_N - u_{N-1}) \approx L \frac{\partial u}{\partial x}$, we get:

$$k_1 L \left. \frac{\partial u}{\partial x} \right|_L + T_a = 0 \tag{12}$$

Defining the macroscopic active stress as $\sigma_{\text{a}} \equiv T_a$ and substituting $GH = k_1 L$, we get the leading edge boundary condition

$$GH \left. \frac{\partial u}{\partial x} \right|_L + \sigma_{\text{a}} = 0 \tag{13}$$

Provided that the $\sigma_a > 0$, there is a negative strain at $x = L$, corresponding to edge retraction.

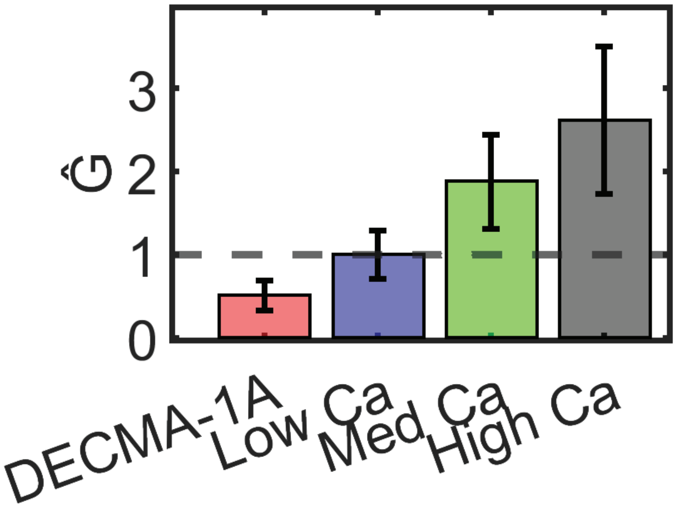

Figure S5: Relative elastic modulus measured with respect to the medium calcium condition derived from the ratio of the penetration length scales squared shows that increasing cell-cell adhesion stiffens the tissue.

#### D Effective elastis modulus of tissues as a function of cell-cell coupling

The variation of penetration length with E-cadherin shown in the main text allowed us to compute the relative elastic moduli of the tissue as a function of cell-cell coupling, measured relative to baseline (medium Ca condition) via the formula $\hat{G} = G_i/G_M = (\lambda_i/\lambda_M)^2$. The subscript $i$ indexes across experimental condition and M corresponds to medium Ca. Inset in Fig. S5 are the experimental results. As cell-cell adhesion increases the effective elastic modulus increases, stiffening the tissue. In contrast, compromising E-cadherin via DECMA-1A softens the tissue.

#### E Non-concentric wound healing assay for finding edge retraction length scale

To place the annular ridge properly in our 2-dimensional wound healing model, we ran an experiment where the wound and the annular ridge were non-concentric and subjected to a single pulse of DC stimulation. The advantage of this assay is that the distance between the ridge and the wound edge, $w$, is continuous. This allowed us to apply stimulation and measure a critical length scale, $w^* \simeq 1.1\,\text{mm}$, representing the minimum distance between the wound edge and the annular ridge field at which does not cause edge retraction. Distances smaller than $w^*$ caused edge significant retraction.

In Fig. S6(A) we show a micrograph of a tissue with the off-centered stimulation region indicated and $w$ labeled. Fig. S6(B) shows the same tissue at $t = 3\,\text{h}$ with $w^*$ labeled. Finally, Fig. S6(C) is a zoomed-in view of the wound region, showing where the edge has curled and where it has maintained integrity. Interestingly, $w^*$ is comparable to the length scale over which information in the orientational order field was propagated in the quasi-1D tissue experiments (Fig. 1F from main

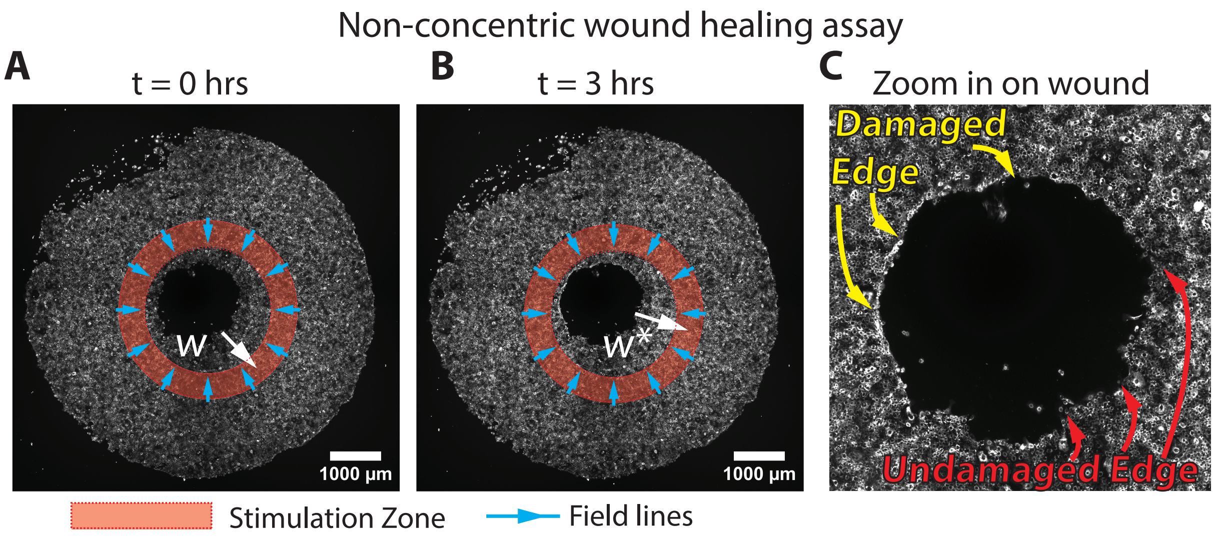

Figure S6: (A) Optical micrograph at $t = 0\,\mathrm{h}$ (pre-stimulation) for the non-concentric wound healing experiment, where the annular ring is off-center with respect to the wound center. This setup yields a continuous distance from the wound edge to the peak of the stimulation region, $w$. (B) Optical micrograph at $t = 3\,\mathrm{h}$ (2.5 hours into stimulation) showing $w^* \simeq 1.1\,\mathrm{mm}$, the critical distance between the wound edge and the stimulation region that prevents edge retraction. (C) Zoomed view of the wound boundary, highlighting regions of damaged and intact edges.

text). Supplemental Video 6 shows this process dynamically. Both nuclear (shown in green) and membrane (shown in red) channels are included. The 3D-printed insert lightly autofluoresces in the nuclear channel, which provides the precise location of the annular ridge.

### F Normalized nuclear density line profiles, healing rates for different stimulation conditions, double pulse kymograph

#### F.1 Normalized nuclear density line profiles

In the main text, we showed spatial heatmaps of the normalized nuclear density for two local stimulation conditions for 2D wound healing. The Continuous Local Stimulation, in which an annular electric field was left on continuously, causing cells to continuously converge on the wound center. This produced a ring of high nuclear density which prevented force and information propagation at the wound edge and quenched wound healing. Conversely, the Single Pulse strategy allowed the nuclear density to relax before jamming occurred. In Fig. S7 we show the corresponding radial density profiles which demonstrate that the Continuous Local Stimulation produces a pronounced peak in nuclear density responsible for the jam (Fig. S7(A)), whereas in the Single Pulse strategy, no density peak is produced (Fig. S7(B)).

#### F.2 Healing dynamics for all 2D strategies

In the main text, many of the healing dynamics were plotted together for compactness and comparison. For clarity, here we include the healing dynamics for each condition plotted separately in Fig. S8. For comparison, we include the unstimulated control cases plotted simultaneously.

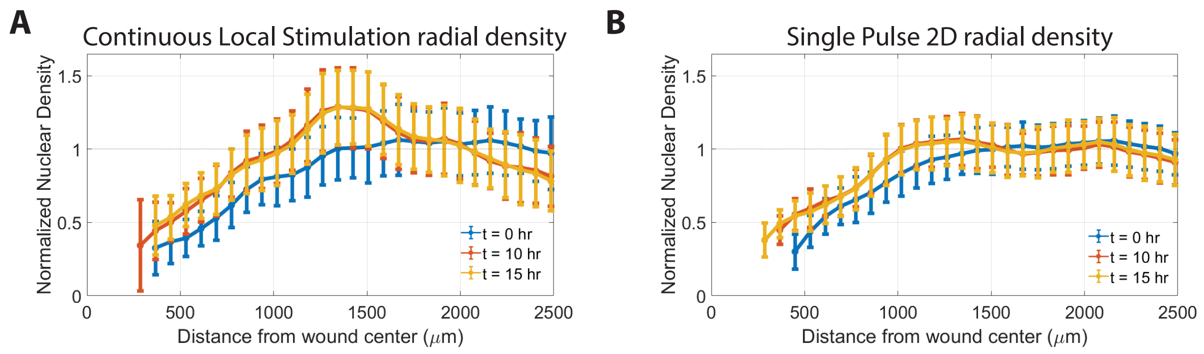

Figure S7: Radial line profiles in time of the normalized nuclear density for (A) Continuous Local Stimulation and (B) Single Pulse. The corresponding normalized density heat maps can be found in the main text. (A) the radial profile for the Continuous Local Stimulation shows a pronounced and stationary peak at $R \simeq 1.5$ mm causing a cellular traffic jam which quenches healing. (B) shows no pronounced peak in normalized density.

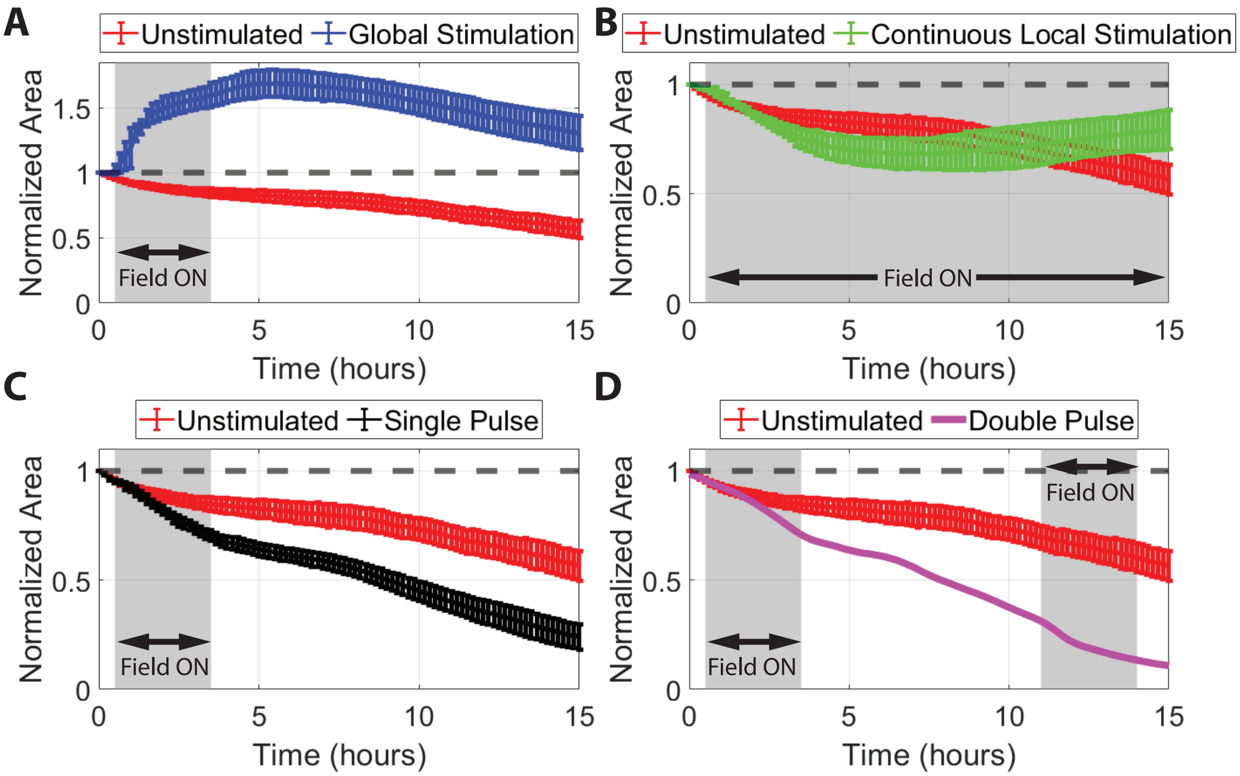

Figure S8: Normalized area dynamics plotted separately for clarity. Each subplot has the unstimulated control for comparison. Included are the dynamics for: (A) Global Stimulation, (B) Continuous Local Stimulation, (C) Single Pulse, and (D) Double Pulse. A dashed line at 1 is included. Note that the $y$-axis range is different for (A).

In addition, we include in Fig. S9 the derivatives of the normalized area, $dA/dt$. Due to the large timestep between optical measurements (10 minutes), there are large error bars on the rates, so they have been omitted for clarity. The horizontal line at $dA/dt = 0$ delineates regions where healing

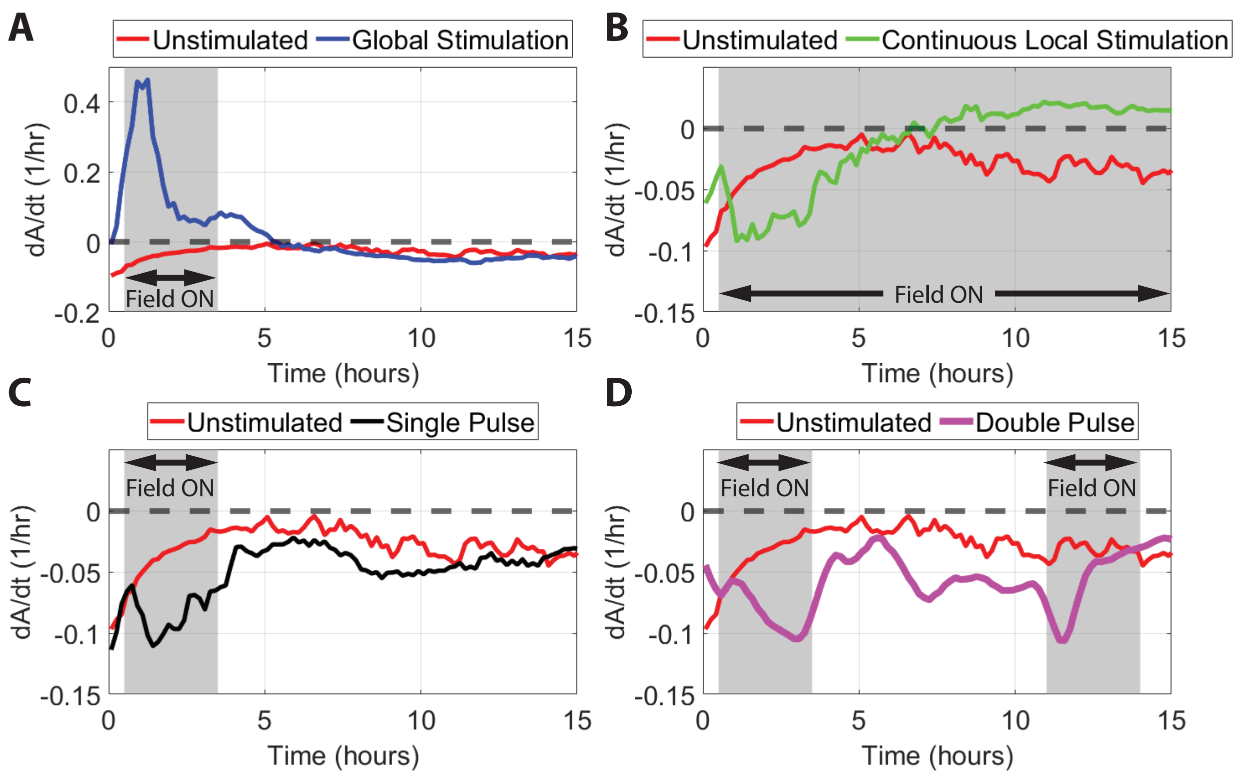

Figure S9: Derivative of unit normalized area dynamics plotted separately. Each subplot has the unstimulated control for comparison. Included are the derivatives for: (A) Global Stimulation, (B) Continuous Local Stimulation, (C) Single Pulse, and (D) Double Pulse. A dashed line at $dA/dt = 0$ separates net positive healing ($dA/dt > 0$) from net negative damage ($dA/dt < 0$). Note that the $y$-axis range is different for (A). Gray zones indicate when the field was on.

rates are positive from those where healing rates are negative. In Fig. S9(A), we observe the Global Stimulation healing rate immediately increase above zero, indicating the wound is expanding in size. We understand this is as a competition between the elasticity of the tissue, which is at minimum a function of the cell-cell coupling, and the applied stimulus. In contrast, for the Continuous Local Stimulation and the Single Pulse Local stimulation plotted in Fig. S9(B) and (C) respectively, we observe that as the field is turned on, there is a near immediate increase in the healing rate from electrotaxis. Finally, for the Double Pulse data shown in Fig. S9(D), we observe two increases in the healing rate commensurate with the stimulation pulses.

##### F.3 Kymograph of Double Pulse stimulation strategy

The Double Pulse control strategy shown in the main text can be compressed to a control kymograph. We show this kymograph in Fig. S10, where two Gaussian pulses centered at radii $R_1 = 1.75$ mm and $R_2 = 1.5$ mm track the wound boundary (purple) in space and are pulsed in time with $\Delta t = 7$ hr delay between pulses (which allows for density relaxation). The kymograph also shows the wound boundary of the unstimulated control (red).

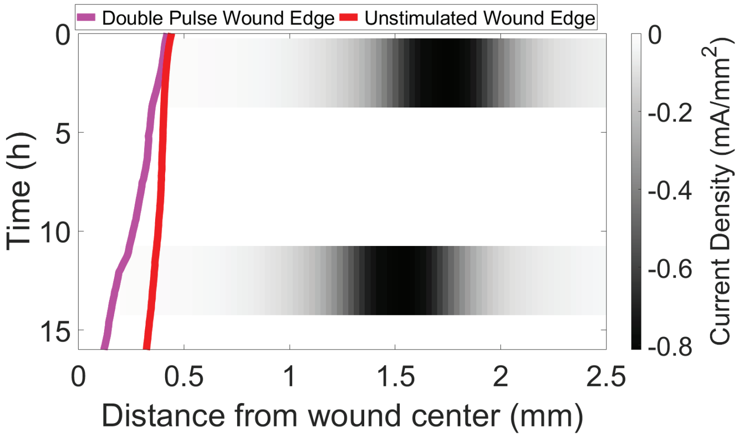

Figure S10: Experimental Double Pulse control kymograph showing two pulses offset in time and space with the effective wound edge is shown in purple along with the unstimulated wound edge in red for comparison.

### G Optimal control for emergence of pulsed strategy: minimally constrained

#### Brief problem description

The foundations and specific instantiations for the optimal control framework are laid out in references [4, 5, 6] with specific instantiations for active systems in [7, 8]. We first start by solving a PDE optimal control problem with minimal constraints to get intuition and then in Section H, we additional physical/experimental constraints to make it more experimentally realizable.

Because the wound is radially symmetric, it can be described in one space dimension, along with the time coordinate. Thus, the wound closure problem can be described with dynamical variables for the tissue density, $\rho(r,t)$, and velocity, $v(r,t)$ on the spatial domain $r \in [0, R]$ and time domain $t \in [0, T_{\text{final}}]$. The tissue density $\rho(r,t)$ is advected by a velocity $v(r,t)$ obtained from a screened (Helmholtz-type) closure driven by a taxis term (density gradients) and by a ring-shaped actuator $u(r,t)$ (model and objective parameters in Table 1). The actuator has *finite agility*: when ON it must dwell for at least a minimum time and cannot exceed a maximum ON time; when OFF it must cool down before it can re-arm; while OFF, its ring center may track a target only at a limited speed (actuator parameters in Table 2). At each time the optimal control solution proposes a candidate ring center in a small window just outside the current rim, guided by an adjoint-based "switching field" $q$; if the signal is strong and stimulation is needed locally, the device turns ON with bounded amplitude. We measure the "open area" by a smooth thresholding of $\rho$ and penalize (i) terminal open area and (ii) unphysical states (creating new wound or overfilling). A terminal adjoint condition encodes the open-area objective, and a shooting iteration adjusts the initial adjoint to satisfy it.

##### Mathematical formulation

The wound–closure problem is formulated as the following constrained optimal control problem:

$$
\begin{aligned}
\min_{(\rho,v,u),\,(A,r_c,z)\in\mathcal{U}_{\text{annulus}}} \quad & J[\rho,u] = \gamma_T A_{\text{open}}[\rho(T_{\text{final}})] + \int_0^{T_{\text{final}}} \Big( B[\rho(t)] + \lambda_u \, \|u(\cdot,t)\|_{L^1_r} \Big)\, dt \\
\text{s.t.} \quad & \partial_t \rho(r,t) + \frac{1}{r}\,\partial_r\big(r\,\rho(r,t)\,v(r,t)\big) = 0, \\
& (I - \nu\,\Delta_r)\,v(r,t) = u(r,t) - \chi\,\partial_r \rho(r,t), \\
& \rho(r,0) = \rho_0(r), \qquad v(0,t) = v(R,t) = 0, \\
& u(r,t) = A(t)\,\mathbf{1}_{\{|r-r_c(t)|\le w_{\text{ring}}/2\}}, \qquad (A,r_c,z)\in\mathcal{U}_{\text{annulus}},
\end{aligned}
\tag{14}
$$

where the radial operator

$$
\Delta_r v := \partial_{rr} v + \frac{1}{r}\,\partial_r v - \frac{1}{r^2} v
$$

matches the screened Helmholtz operator used in the model (see Table 1 for parameter values). The running barrier functional $B[\rho(t)]$ penalizes undesirable density profiles (reopening of tissue that has once healed and overfilling beyond a biologically plausible upper bound), while the terminal open–area metric $A_{\text{open}}[\rho(T_{\text{final}})]$ and the radial $L^1$–norm of the control are given by

$$
B[\rho(t)] = \int_0^R \Big( \kappa_{\text{low}}\, H_+\big(\rho_{\text{thr}} - \rho(r,t)\big)\, S(r,t) + \kappa_{\text{high}}\, H_+\big(\rho(r,t) - \rho_{\max}\big) \Big)\, 2\pi r\, dr, \tag{15}
$$

$$
A_{\text{open}}[\rho(T_{\text{final}})] = \frac{2\pi}{\pi R^2} \int_0^R \sigma\big(a_{\text{thr}}(\rho_{\text{thr}} - \rho(r,T_{\text{final}}))\big)\, r\, dr, \tag{16}
$$

$$
\|u(\cdot,t)\|_{L^1_r} := \int_0^R |u(r,t)|\, 2\pi r\, dr. \tag{17}
$$

Here $H_+$ is a smooth hinge (softplus) approximation of $x \mapsto \max(0,x)$, $\sigma(z) = 1/(1+e^{-z})$ is a logistic function, and $S(r,t)$ is a dynamic mask that records locations that have ever crossed the healing threshold $\rho_{\text{thr}}$. Thus $B[\rho(t)]$ acts as a soft state constraint, discouraging both the re-creation of wound in regions that once healed (through $S$) and nonphysical overfilling beyond $\rho_{\max}$ (parameters in Table 4). The radial $L^1$-penalty $\lambda_u \|u(\cdot,t)\|_{L^1_r}$ promotes temporally sparse, spatially localized actuation and, together with the pointwise bound $|u| \le U_{\max}$, implies via Pontryagin's minimum principle a bang–bang structure for the optimal annulus amplitude,

$$
A(t) = \pm U_{\max}\, \mathrm{sign}\big(-q(r_c(t),t)\big),
$$

with an effective switching condition $|q(r_c(t),t)| > \lambda_{\text{ring}}$ (see Table 2).

The admissible–control set $\mathcal{U}_{\text{annulus}}$ encodes the geometry of the annulus and finite–agility constraints (dwell and cooldown times, bounded tracking speed for $r_c(t)$, and an activation threshold on the switching field), summarized in Table 2.

**State, actuator, mask, and open-area metric.** The state variable is $\rho(r,t) \ge 0$. A dynamic "once healthy, always healthy" mask $S(r,t) \in \{0,1\}$ records locations that have ever exceeded a healing threshold $\rho_{\text{thr}}$:

$$
S(r,t^+) \;=\; \max\big(S(r,t^-),\, \mathbf{1}_{\{\rho(r,t)>\rho_{\text{thr}}\}}\big).
$$

The actuator is a top-hat annulus of width $w_{\text{ring}}$ centered at $r_c(t)$ with held amplitude $A(t)$ while ON:

$$u(r,t) \;=\; A(t)\,\mathbf{1}_{\{|r-r_c(t)|\le \frac{1}{2}w_{\text{ring}}\}}.$$

A smooth open-area fraction is computed via a logistic threshold with sharpness $a_{\text{thr}}$,

$$\mathcal{A}\big(\rho(\cdot,t)\big) = \frac{1}{\pi R^2}\int_0^R \sigma\big(a_{\text{thr}}(\rho_{\text{thr}}-\rho(r,t))\big)\,(2\pi r)\,dr, \qquad \sigma(z)=\frac{1}{1+e^{-z}},$$

with terminal variational gradient

$$\frac{\delta\mathcal{A}}{\delta\rho}(r,T_{\text{final}}) = -\frac{2\pi r}{\pi R^2}\,a_{\text{thr}}\,\sigma(z)\big(1-\sigma(z)\big)\Big|_{z=a_{\text{thr}}(\rho_{\text{thr}}-\rho(r,T_{\text{final}}))},$$

and associated parameter values in Tables 1 and 4.

**Transport of $\rho$ and screened closure for $v$.** Density is transported conservatively by $v$:

$$\partial_t\rho \;+\; \frac{1}{r}\,\partial_r\big(r\,\rho\,v\big) \;=\; 0, \qquad (r,t)\in(0,R)\times(0,T_{\text{final}}). \tag{18}$$

The velocity is obtained from a screened elliptic relation (with the radial Laplacian $L=\Delta_r$):

$$\big(I-\nu\Delta_r\big)\,v \;=\; u \;-\; \chi\,\partial_r\rho, \qquad v(0,t)=0, \quad v(R,t)=0. \tag{19}$$

Here, $\nu>0$ is the smoothing and $\chi\ge 0$ the taxis sensitivity (see Table 1).

**Adjoint transport, terminal data, and switching field.** Let $\phi(r,t)$ be the adjoint variable associated with $\rho$. It is transported forward with $-v$ and driven by smooth barrier sources that discourage reopening or overfilling:

$$\partial_t\phi + v\,\partial_r\phi \;=\; -\,\mu(\rho,S), \qquad \phi(r,T_{\text{final}}) \;=\; \gamma_T\,\frac{\delta\mathcal{A}}{\delta\rho}(r,T_{\text{final}}), \tag{20}$$

with

$$\mu(\rho,S) = \kappa_{\text{low}}\ \text{softplus}\Big(\frac{\rho_{\text{thr}}-\rho}{\tau_{\text{hinge}}}\Big)\,S \;+\; \kappa_{\text{high}}\ \text{softplus}\Big(\frac{\rho-\rho_{\max}}{\tau_{\text{hinge}}}\Big), \quad \text{softplus}(x)=\log(1+e^x),$$

and parameters as in Table 4. A *switching field* $q(r,t)$ is obtained from the same screened operator and the adjoint sensitivity,

$$\big(I-\nu\Delta_r\big)\,q \;=\; S_q\,\rho\,\partial_r\phi, \qquad S_q\in\{\pm 1\}, \tag{21}$$

and is used to locate and evaluate candidate centers via $|q|$ (with sign $S_q$ in Table 4).

**Finite-agility actuator logic.** Let $z(t)\in\{0,1\}$ be ON/OFF, with ON and OFF clocks, a held amplitude $A(t)$, and center $r_c(t)$. At each time:

- Define a rim location $r_w(t)$ as the smallest $r$ with $\rho(r,t)\ge\rho_{\text{thr}}$. Restrict candidates $r^\star$ to the window $[\,r_w+b_{\text{in}},\, r_w+b_{\text{out}}\,]$ (or search globally if empty) and take the maximizer of $|q(\cdot,t)|$.
- *ON branch* ($z=1$): hold $r_c$ and $A$; turn OFF if the minimum dwell has elapsed and local need has ended, or if the hard ON cap is reached.

- *OFF branch* ($z = 0$): set $A = 0$; move $r_c$ toward $r^\star$ with bounded speed $|\dot r_c| \le v_{\text{track}}$; re-arm (turn ON) only if cooldown elapsed, $|q(r^\star, t)| > \lambda_{\text{ring}}$, and the ring overlaps a sub-threshold region.
- On switching ON: set $A = \pm U_{\max}\,\mathrm{sign}\big(-q(r^\star, t)\big)$, $r_c \leftarrow r^\star$, reset clocks.

All actuator parameters and constraints are collected in Table 2.

**Terminal objective and shooting.** The scalar objective is the terminal open-area fraction $\mathcal{A}(\rho(\cdot, T_{\text{final}}))$ (encoded via the terminal adjoint condition in (20)). A shooting iteration adjusts $\phi(r, 0)$ so that the terminal adjoint $\phi(r, T_{\text{final}})$ matches $\gamma_T\, \delta\mathcal{A}/\delta\rho$.

##### Numerical implementation

**Grids and operators.** Uniform grids: $r_i = i\,\Delta r$ on $[0, R]$ and $t^n = n\,\Delta t$ on $[0, T_{\text{final}}]$. Spatial derivatives use centered differences; the radial Laplacian is $Lf \approx f_{rr} + \frac{1}{r} f_r$ with regularized $1/r$ at $r = 0$. The screened solves (19), (21) are tridiagonal and evaluated with a Thomas algorithm; boundary rows enforce $v(0) = v(R) = 0$. The main discretization and solver constants are summarized in Table 3.

**Conservative advection for $\rho$.** We discretize (18) in finite-volume form to respect cylindrical geometry:

$$\frac{\rho_i^{n+1} - \rho_i^n}{\Delta t} = -\frac{1}{r_i}\,\frac{(rF)^n_{i+\frac{1}{2}} - (rF)^n_{i-\frac{1}{2}}}{\Delta r}, \qquad F = \rho\, v,$$

with upwinded face fluxes and zero outer flux via $v(R, t) = 0$. To maintain a target CFL, we split $\Delta t$ into $N_{\text{sub}} = \max\big\{2,\ \lceil \|v\|_\infty\, \Delta t/(C_{\text{CFL}} \Delta r)\rceil\big\}$ substeps ($C_{\text{CFL}} \approx 0.6$).

**Semi-Lagrangian adjoint transport and barriers.** Equation (20) is advanced by semi-Lagrangian back-tracking with clipped linear interpolation and an explicit sink $-\mu\,\Delta t$. After each $\rho$ update we refresh $S^{n+1} = \max\big(S^n, \mathbf{1}_{\{\rho^{n+1} > \rho_{\text{thr}}\}}\big)$, then construct $\mu(\rho^{n+1}, S^{n+1})$ using the parameters in Table 4.

**Actuator evaluation per step.** At each $t^n$: (i) solve (19) for $v^n$; (ii) advance $\rho$ with the FV step (with substeps if needed); (iii) update $S$ and step $\phi$; (iv) solve (21) for $q^n$; (v) run the finite-agility state machine to produce $u^{n+1}$ (Tables 2 and 3).

**Shooting update.** Define the terminal residual $r_T(r) = \phi(r, T_{\text{final}}) - \gamma_T \frac{\delta \mathcal{A}}{\delta \rho}(r, T_{\text{final}})$. Transport $r_T$ backward through the recorded velocity history to $t = 0$ to get $r_0$, then update $\phi_0 \leftarrow \phi_0 - \eta\, r_0$ (small step $\eta$), and repeat.

#### H Experimentally constrained optimal control: Brief problem description

We now proceed onto the PDE optimal control problem with additional physical/experimental constraints to make it more realistic. As in the previous section, we study a radially symmetric model of wound closure on a disk of radius $R$ over a time horizon $[0, T]$. The tissue is represented by a density field $\rho(r, t)$ that is transported by a radial velocity $v(r, t)$ and diffuses on the healed

Table 1: Model, closure, and objective parameters.

| Symbol / name | Description | Value (units) |
|---|---|---|
| $R$ | Domain radius | 5.0 mm |
| $T_{\text{final}}$ | Time horizon | 20.0 h |
| $r_{w0}$ | Initial rim location | 1.2 mm |
| $\rho_{\max}$ | Density upper cap | 4.0 |
| $\rho_{\text{thr}}$ | Healing threshold | $0.25\,\rho_{\max}$ |
| $\chi$ | Taxis sensitivity | 0.010 |
| $\nu$ | Screening coefficient | 0.2 |
| $U_{\max}$ | Control amplitude cap | 0.10 |
| $a_{\text{thr}}$ | Open-area logistic sharpness | 8.0 |
| $\gamma_T$ | Terminal weight (open area) | 10.0 |
| $S_q$ | Sign in $(\nu L - I)q = S_q\,\rho\,\partial_r\phi$ | $-1.0$ |
| BCs | Velocity boundary conditions | $v(0,t) = v(R,t) = 0$ |

Table 2: Actuator geometry and finite-agility logic.

| Symbol / name | Description | Value (units) |
|---|---|---|
| $w_{\text{ring}}$ | Annulus width | 0.50 mm |
| $b_{\text{in}}$ | Candidate window start beyond rim | 0.10 mm |
| $b_{\text{out}}$ | Candidate window end beyond rim | 0.70 mm |
| $\lambda_{\text{ring}}$ | ON threshold on $\lvert q\rvert$ at center | $10^{-3}$ |
| $\tau_{\text{on}}^{\min}$ | Minimum uninterrupted ON (dwell) | 0 h |
| $\tau_{\text{on}}^{\max}$ | Maximum ON time (hard cap) | $0.1\,T_{\text{final}}$ |
| $\tau_{\text{off}}^{\min}$ | Required OFF time (cooldown) | $0.01\,T_{\text{final}}$ |
| $v_{\text{track}}$ | Max center tracking speed while OFF | 0.50 mm/h |
| Switching center | Argmax of $\lvert q\rvert$ in $[r_w+b_{\text{in}},\ r_w+b_{\text{out}}]$ | — |
| Held amplitude | While ON, $A(t) = \pm U_{\max}\,\text{sign}(-q)$ | — |

Table 3: Discretization and numerical constants.

| Symbol / name | Description | Value (units) |
|---|---|---|
| $N_r$ | Radial grid points | 512 |
| $N_t$ | Time steps | 1500 |
| $\Delta r$ | Radial spacing | $R/(N_r-1)$ |
| $\Delta t$ | Time step | $T_{\text{final}}/(N_t-1)$ |
| Advection of $\rho$ | Finite-volume, geometric flux form | FV |
| Adjoint transport | Semi-Lagrangian + explicit source | SL |
| Linear solves | Tridiagonal Thomas solver | — |
| EPS | Small numerical guard | $10^{-12}$ |

tissue. The wound edge $r_w(t)$ is the radius where the tissue transitions from "open" to "healed" and moves kinematically with the local tissue velocity. We accelerate closure using an annular (ring-shaped) actuator that applies a spatially localized control $u(r,t)$ in short pulses. When a pulse turns ON, its ring center is frozen at a prescribed standoff outside the then-current edge; between pulses the actuator is OFF. The optimization variables are the pulse start times and the standoffs. The objective is to minimize the final wound size while enforcing soft safety constraints and solution smoothness.

Table 4: Adjoint, barriers, and open-area metric.

| Symbol / name | Description | Value (units) |
|---|---|---|
| $\kappa_{\text{low}}$ | Barrier: penalize new wound (on dynamic mask) | 3.0 |
| $\kappa_{\text{high}}$ | Barrier: penalize overfill | 15.0 |
| $\tau_{\text{hinge}}$ | Hinge softness (density units) | 0.05 |
| Open area $\mathcal{A}$ | Logistic on $(\rho_{\text{thr}} - \rho)$, normalized by $\pi R^2$ | — |
| $\delta\mathcal{A}/\delta\rho$ | Terminal gradient for adjoint BC | from $a_{\text{thr}}$ |
| $q$-equation | $(\nu L - \alpha I)q = S_q\,\rho\,\partial_r\phi$ | $S_q = -1$ |

### Mathematical formulation

**State, control, and masks.** The state is $(\rho, r_w)$ with $\rho(r,t) \geq 0$, $r \in [0,R]$, $t \in [0,T]$. Two smooth masks separate healed tissue from void: a tissue mask

$$m(r,t) = \frac{1}{1 + e^{-\kappa_m\,(r - r_w(t))}}$$

and a void guard

$$g(r,t) = \frac{1}{1 + e^{-\kappa_v\,(r - r_w(t))}}\;\frac{1}{1 + e^{-\kappa_v\,(r - r_g(t))}}, \qquad r_g(t) = \max\{\, r_w(t) - n_g\,\Delta r,\; 0\,\}.$$

We use the effective mask $m_{\text{eff}}(r,t) = m(r,t)\,g(r,t)$ so that transport and integrals are restricted to tissue.

**Dynamics and kinematics.** Density evolves by advection–diffusion on the masked tissue; the velocity is obtained from a screened (Helmholtz) closure forced by taxis (density gradients) and control. The edge evolves kinematically with the local velocity sampled around the edge. Writing the radial operators $\partial_r f = f_r$ and $\Delta_r f = f_{rr} + \frac{1}{r} f_r$, we use

$$\partial_t \rho \;+\; \frac{1}{r}\,\partial_r\big(r\,\rho\,v\big) = D\,\Delta_r\big(\rho\,g\big) \qquad \text{on } (0,R)\times(0,T), \tag{22}$$

$$\big(I - \nu\,\Delta_r\big)\,v = -\,\chi\,\partial_r \rho \;+\; u(r,t) \qquad \text{on } (0,R)\times(0,T), \tag{23}$$

$$\dot r_w(t) = \big\langle v(\,\cdot\,,t)\big\rangle_{\text{edge}}, \tag{24}$$

with regularity at $r = 0$ and Neumann-type behavior at $r = R$ in the discretization. The edge average $\langle v\rangle_{\text{edge}}$ is a narrow Gaussian-weighted average of $v$ around $r = r_w(t)$. Initial data are $\rho(r,0) = \rho_0(r)$ (a smooth ramp across $r_w(0)$) and $r_w(0) = r_{w0}$.

**Control pulses and ring construction.** We parameterize $P$ pulses by start times $t_{\text{on}}^{(p)}$ and standoffs $d^{(p)} \in [S_{\min}, L_{\text{prop}}]$, mapped from unconstrained variables via sigmoids. Each pulse has duration $\tau$ and a smooth top-hat $W^{(p)}(t)$ that is *cropped* to the ON window (no tails outside $[t_{\text{on}}^{(p)}, t_{\text{on}}^{(p)} + \tau]$). When a pulse turns ON, its center freezes at

$$r_c^{(p)} \;=\; r_w\big(t_{\text{on}}^{(p)}\big) + d^{(p)}.$$

The spatial ring is a normalized Gaussian of width $\sigma_r$ restricted to tissue:

$$k^{(p)}(r,t) \;=\; \frac{\exp\!\big(-\tfrac{1}{2}\tfrac{(r - r_c^{(p)})^2}{\sigma_r^2}\big)\,m(r,t)}{\displaystyle\int_0^R \exp\!\big(-\tfrac{1}{2}\tfrac{(s - r_c^{(p)})^2}{\sigma_r^2}\big)\,m(s,t)\,s\,ds + \varepsilon},$$

and the control is a signed sum with amplitude $J_0$,

$$u(r,t) \;=\; -\,J_0 \sum_{p=1}^{P} k^{(p)}(r,t)\, W^{(p)}(t)\, \mathbf{1}_{[t_{\mathrm{on}}^{(p)},\, t_{\mathrm{on}}^{(p)}+\tau]}(t).$$

**Objective: terminal goal and running penalties.** We minimize a weighted sum that encodes a small final wound, safety, mass consistency, damping when OFF, and smoothness of $\rho$ and $u$:

$$\begin{aligned}
\mathcal{J} = \;& \underbrace{L_{\mathrm{wound}}\, r_w(T)^2}_{\text{final wound size (smaller is better)}} \;+\; \underbrace{L_{\mathrm{edge}} \int_0^T \big[\dot r_w(t)\big]_+^2\, dt}_{\text{penalize outward edge motion over time}} \;+\; \underbrace{L_{\mathrm{mass}} \int_0^T \big(M(t) - M_0\big)^2\, dt}_{\text{preserve tissue mass on the healed region}} \\
&+ \underbrace{L_{\mathrm{safe}} \int_0^T \sum_{p=1}^{P} \big[S_{\mathrm{safe}} - (r_c^{(p)} - r_w(t))\big]_+^2\, \mathbf{1}_{\mathrm{ON}}^{(p)}(t)\, dt}_{\text{enforce minimum ring–edge standoff while a pulse is ON}} + \underbrace{L_{\mathrm{jam}} \int_0^T \exp\Big(-\frac{1}{2}\frac{(r_c^{(2)} - r_c^{(1)})^2}{\sigma_{\mathrm{jam}}^2}\Big)\, dt}_{\text{avoid ring jamming (keep centers separated)}} \\
&+ \underbrace{L_{\mathrm{damp}} \int_0^T \Big(\int_0^R v(r,t)^2\, m_{\mathrm{eff}}(r,t)\, r\, dr\Big)\, \mathbf{1}_{\text{all OFF}}(t)\, dt}_{\text{damp residual motion when all pulses are OFF}} \\
&+ \underbrace{\iint \big(L_{\rho r}\, |\partial_r \rho|^2 + L_{\rho t}\, |\partial_t \rho|^2\big)\, m_{\mathrm{eff}}\, r\, dr\, dt}_{\text{regularize density: smooth in space and time}} + \underbrace{\iint \big(L_{ur}\, |\partial_r u|^2 + L_{ut}\, |\partial_t u|^2\big)\, m_{\mathrm{eff}}\, r\, dr\, dt}_{\text{regularize control: smooth in space and time}} \\
&+ \underbrace{L_{\mathrm{ton}}\ \mathrm{dist}\big(t_{\mathrm{on}}^{(1)}, [T_1^{\min}, T_1^{\max}]\big)^2}_{\text{encourage first pulse to start within a target time window}} . \qquad (25)
\end{aligned}$$

Here $[x]_+ = \max\{x, 0\}$ is implemented by a smooth hinge; $M(t) = \int_0^R \rho\, m_{\mathrm{eff}}\, r\, dr$ is tissue mass with baseline $M_0$; indicators are smooth surrogates in code.

**Optimal control problem** *Find pulse start times and standoffs* $\{t_{on}^{(p)}, d^{(p)}\}_{p=1}^{P}$ *that minimize* $\mathcal{J}$ *in (25), subject to the PDEs (22)–(23) and the edge law (24), the control construction (frozen centers while ON; cropped windows), and initial data* $\rho(r,0) = \rho_0(r)$, $r_w(0) = r_{w0}$.

### Hyperparameters

Tables 5–6 list physical/model and numerical/optimization hyperparameters used in our experiments.

### Numerical implementation

**Grids and operators.** We use uniform grids: $r_i = i\,\Delta r$ for $i = 0, \ldots, N_r - 1$ on $[0, R]$, and $t^n = n\,\Delta t$ for $n = 0, \ldots, N_t$ on $[0, T]$. Radial derivatives use centered differences with "edge padding" at $i = 0$ and $i = N_r - 1$ to impose regularity (at $r = 0$) and Neumann-type behavior (at $r = R$). The geometric divergence is discretized in conservative form:

$$\nabla_r \cdot (rF)/r \;\approx\; \frac{1}{r_i}\, \frac{(rF)_{i+\frac{1}{2}} - (rF)_{i-\frac{1}{2}}}{\Delta r},$$

with upwinded face fluxes for $F = \rho v$ to improve stability of advection.

Table 5: Physical and model parameters.

| Symbol / name | Description | Value (units) |
|---|---|---|
| $R$ | Domain radius | 5.0 mm |
| $T$ | Time horizon | 20.0 h |
| $D$ | Diffusivity | $0.02\,\mathrm{mm}^2/\mathrm{h}$ |
| $\chi$ | Taxis coefficient | 0.25 |
| $\nu$ | Screening (Helmholtz) coefficient | 0.15 |
| $P$ | Number of control pulses | 2 |
| $\tau$ | Pulse duration | 3.0 h |
| $[G_{\min}, G_{\max}]$ | Gap range between pulses | [4.0, 5.0] h |
| $J_0$ | Control amplitude scale | 0.9 |
| $\sigma_r$ | Ring width | 0.175 mm |
| $S_{\min}$ | Min standoff (mapping lower bound) | 0.3 mm |
| $L_{\text{prop}}$ | Max standoff (mapping upper bound) | 1.0 mm |
| $S_{\text{safe}}$ | Safety standoff during ON | 0.4 mm |
| $\beta_{\text{safe}}$ | Safety hinge sharpness | 10.0 |
| $\sigma_{\text{jam}}$ | Jamming proximity scale | 0.25 mm |
| $K_{\text{mask}}$ | Tissue-mask sharpness | 6.0 |
| $K_{\text{void}}$ | Void-guard sharpness | 120.0 |
| $n_g$ | Guard band (cells) | 20 |
| $\rho_{\min}, \rho_{\max}$ | Density clipping | $10^{-8}$, 10 |
| $L_{\text{wound}}$ | Final wound weight | $10^4$ |
| $L_{\text{edge}}$ | Outward edge speed weight | 1.0 |
| $L_{\text{mass}}$ | Mass consistency weight | 10.0 |
| $L_{\text{safe}}$ | Safety weight | 1.0 |
| $L_{\text{jam}}$ | Jamming weight | 0.2 |
| $L_{\text{damp}}$ | Damping-when-OFF weight | 1.0 |
| $L_{\rho r}, L_{\rho t}$ | $\rho$ spatial/temporal regularization | 1.0, 1.0 |
| $L_{ur}, L_{ut}$ | $u$ spatial/temporal regularization | 1.0, 1.0 |
| $[T_1^{\min}, T_1^{\max}]$ | First-pulse timing window | [0.0, 1.0] h |
| $L_{\text{ton}}$ | First-pulse timing weight | $10^3$ |

**Velocity solve** At each time step we solve $(I - \nu\Delta_r)v = -\chi\,\partial_r\rho + u$ with a tridiagonal (Thomas) solver. Boundary rows implement $r{=}0$ regularity and $r{=}R$ Neumann-like behavior. The right-hand side is guarded against NaNs and small denominators.

**Time stepping for $\rho$ and mass correction.** We update $\rho$ explicitly on tissue using

$$\rho^{n+1} \;=\; \rho^n \;+\; \Delta t\,\Big(D\,\Delta_r(\rho^n g^n) \;-\; \nabla_r\cdot(\rho^n v^n)\Big)\,m^n_{\text{eff}},$$

then apply a multiplicative mass correction on tissue to keep $M^{n+1} \approx M_0$:

$$\rho^{n+1} \;\leftarrow\; \rho^{n+1}\Big(1 + \lambda_{\text{mc}}\,(M_0/M^{n+1} - 1)\,m^n_{\text{eff}}\Big),$$

followed by clipping to $[\rho_{\min}, \rho_{\max}]$ and reapplying the guard. A narrow Gaussian weight around $r^n_w$ yields $\dot r^n_w = \sum_i w_i(r^n_w)\,v^n_i$ and $r^{n+1}_w = r^n_w + \Delta t\,\dot r^n_w$.

Table 6: Numerical and optimization parameters.

| Symbol / name | Description | Value (units) |
|---|---|---|
| $N_r$ | Radial grid size | 192 |
| $N_t$ | Time steps | 1600 |
| $\Delta r$ | Radial spacing | $R/(N_r - 1)$ |
| $\Delta t$ | Time step | $T/N_t$ |
| $\varepsilon$ | Normalization guard (denominator) | $10^{-10}$ |
| $\lambda_{\mathrm{mc}}$ | Mass-correction relaxation | 1.0 |
| $\varepsilon_{\mathrm{mask}}$ | Small mask constant | $10^{-8}$ |
| $\sigma_J$ | Edge-sampling width | $\max(2\Delta r,\, 2R/N_r)$ |
| Optimizer | Adam learning rate, steps | 0.05, 800 iterations |
| Param map | Sigmoid maps for $t_{\mathrm{on}}, d$ | as in text |
| BCs | $r$=0 regularity; $r$=$R$ Neumann-like | tridiagonal Helmholtz |

**Control construction and ON/OFF logic.** At time $t^n$, each pulse uses a smooth top-hat $W^{(p)}(t^n)$ that is multiplied by a Boolean ON window to crop tails outside $[t_{\mathrm{on}}^{(p)}, t_{\mathrm{on}}^{(p)} + \tau]$. When a pulse transitions OFF→ON, we freeze its center $r_c^{(p)} \leftarrow r_w^n + d^{(p)}$; while ON, $r_c^{(p)}$ remains fixed. The instantaneous control is

$$u_i^n \;=\; -\,J_0 \sum_{p=1}^{P} \frac{\exp\!\big(-\tfrac{1}{2}\tfrac{(r_i - r_c^{(p)})^2}{\sigma_r^2}\big)\, m_i^n}{\sum_j \exp\!\big(-\tfrac{1}{2}\tfrac{(r_j - r_c^{(p)})^2}{\sigma_r^2}\big)\, m_j^n\, r_j\, \Delta r + \varepsilon}\; W^{(p)}(t^n)\, \mathbf{1}_{\mathrm{ON}}^{(p)}(t^n).$$

**Regularization, safety, and stabilization.** All inequality-type requirements (minimum standoff, outward edge motion, timing windows) are enforced with smooth softplus/hinge penalties to preserve differentiability. The safety standoff acts only while a pulse is ON. A damping term penalizes $\int v^2$ when all pulses are OFF to avoid spurious motion. Clipping of $\rho$ prevents blow-up and negative densities; small guards ($\varepsilon$) avoid division by zero.

**Optimization.** Pulse parameters are mapped via sigmoids to enforce bounds and minimum gaps. The objective $\mathcal{J}$ is evaluated by Riemann sums ($r_i \Delta r$ in space and $\Delta t$ in time). We differentiate through the full time-stepping loop by reverse-mode automatic differentiation and update parameters with Adam (learning rate 0.05, 800 iterations in the reported runs).

**Practical CFL guidance.** Although masks and smoothing improve robustness, stability is aided by choosing $\Delta t$ to respect diffusion and advection scales, e.g. $D\,\Delta t/\Delta r^2 \lesssim \mathcal{O}(0.5)$ and $\max_i |v_i|\,\Delta t/\Delta r \lesssim \mathcal{O}(0.5)$ in typical runs.

### I Description of Movies

1. Representative timelapse video of "Unstimulated" 2D wound with no applied electric field to show baseline healing rates.

2. Representative timelapse video of 2D wound with "Global Stimulation" control with monopolar spatial stimulation delivered in a single 3-hour pulse.

3. Timelapse video of a quasi-1D (6 mm × 1.5 mm tissue of medium Ca keratinocytes in a homogeneous 2.5 V/cm electric field. Edge retracts from the global command.

4. Timelapse video of quasi-1D (10 mm × 1.5 mm) tissue of medium Ca keratinocytes locally stimulated at the center shows localized migration

5. Average heatmap videos of speed and orientational order in response to local stimulation shows localized response in speed and broadened response in orientational order.

6. Representative timelapse video of non-concentric wound healing experiments showing the nuclear channel (green) and cytoplasmic membrane channel (red), highlighting regions where edge retraction occurs.

7. Representative timelapse video of 2D wound with "Continuous Local Stimulation" with stimulation on from $t > 0.5$ h onward

8. Representative timelapse video of 2D wound "Single Pulse" delivering local stimulation in a single 3-hour pulse.

9. Average nuclear density heatmap videos of "Continuous Local Stimulation" and "Single Pulse" showing density build-up in the "Continuous Local Stimulation" experiments.

10. Experimental double pulse sequence where both pulses are at the same location in space, $R_1 = R_2 = 1.75$ mm showing that edge tracking is critical.

11. Top: healing dynamics of $\rho(r, t)$ and $u(r, t)$ without control. Bottom: healing dynamics of $\rho(r, t)$ and $u(r, t)$ for experimentally constrained optimal control solution.

12. Representative timelapse video of 2D wound "Double Pulse" delivering local stimulation at $R_1 = 1.75$ mm and $R_2 = 1.25$ mm in two 3-hour pulses.

13. Experimental double pulse sequence where $R_1 = 1.75$ mm, $R_2 = 0.75$ mm. The second pulse is too close to the wound edge and generates edge retraction.

[5] Arthur Earl Bryson. *Applied optimal control: optimization, estimation and control*. Routledge, 2018.

[6] Lev Semenovich Pontryagin. *Mathematical theory of optimal processes*. Routledge, 2018.

[7] Sumit Sinha, Vishaal Krishnan, and L Mahadevan. Optimal control of interacting active particles on complex landscapes. *arXiv preprint arXiv:2311.17039*, 2023.

[8] Vishaal Krishnan, Sumit Sinha, and L Mahadevan. Hamiltonian bridge: A physics-driven generative framework for targeted pattern control. *arXiv preprint arXiv:2410.12665*, 2024.
